# Revisiting the Kuleshov effect with authentic films: A behavioral and fMRI study

**DOI:** 10.1101/2023.12.04.569135

**Authors:** Zhengcao Cao, Yashu Wang, Liangyu Wu, Yapei Xie, Zhichen Shi, Yiren Zhong, Yiwen Wang

## Abstract

As a fundamental film theory, montage theory posits that film editing significantly influences the viewer’s perception. The Kuleshov effect, a central concept in montage theory, proposes that the emotional interpretation of neutral facial expressions is influenced by the accompanying emotional scene in a face-scene-face sequence. However, concerns persist regarding the validity of previous studies, often employing inauthentic film materials like static images, leaving the question of its existence in authentic films unanswered. This study addresses these concerns by utilizing authentic films in two experiments. In Experiment 1, multiple film clips were captured under the guidance of a professional film director and seamlessly integrated into authentic film sequences. A total of 59 participants viewed these face-scene-face film sequences and were tasked with rating the valence of neutral faces. The findings revealed that the interpretation of emotion in neutral faces is significantly influenced by the accompanying fearful or happy scene, eliciting perceptions of negative or positive emotion from the neutral face. These results affirm the existence of the Kuleshov effect within authentic film sequences. In Experiment 2, 31 participants engaged in a similar task under functional magnetic resonance imaging (fMRI). The results revealed neural correlates supporting the existence of the Kuleshov effect at a neural level. These correlates include the cuneus, precuneus, hippocampus, parahippocampal gyrus, post cingulate gyrus, orbitofrontal cortex, fusiform gyrus, and insula. These findings also underscore the contextual framing inherent in the Kuleshov effect. By seamlessly integrating film theory and cognitive neuroscience experiments, this comprehensive study provides robust evidence supporting the existence of the Kuleshov effect. It significantly enhances our understanding of film editing and its profound impact on viewers’ perception.

## 1. Introduction

In film theory, montage entails the combination of two or more images through film editing, creating new meanings from the foundational frames (Bordwell, 1972). From a psychological perspective, film editing significantly influences the viewer’s perception of the movie, allowing them to experience emotions beyond those portrayed by the actors (Calbi et al., 2017; Carroll & Carroll, 1993; Tan, 2018). The Kuleshov effect, a renowned example of montage, aims to investigate how film editing shapes the viewer’s interpretation of emotions (Barratt et al., 2016; Prince & Hensley, 1992).

The Kuleshov effect was detected through the Kuleshov experiment, which consists of a series of frames assembled in a face-scene-face sequence (Pudovkin, 1970). This sequence includes segments featuring an actor with a neutral expression, followed by an emotional scene, and concluding with a repetition of the actor displaying a neutral expression. In this experiment, viewers were tasked with assessing the emotions portrayed by the actor’s face. Despite the actor’s facial expression remaining consistent, the viewer frequently attributed different emotional states to these actors, and this emotional state is intricately linked to the accompanying emotional scene. For example, when the emotional scene depicts a dead girl lying in a cabin, the viewer perceives fear in the neutral face. However, when the emotional scene features an alluring woman reclining on a couch, the viewer interprets happiness in the neutral face.

Since Kuleshov’s initial pilot experiment on the Kuleshov effect in the 1920s (Pudovkin, 1970), references to the Kuleshov effect in film textbooks suggest that the effect is well established (McDonald, 2022; Reisz & Millar, 1971). However, due to the vagueness and lack of control in the initial experiment, it is not clear whether the Kuleshov effect truly exists. To address this question, numerous scientific experiments have attempted to replicate the study, utilizing diverse experimental materials and paradigms throughout the past century (Barratt et al., 2016; Ekman, 1971; Prince & Hensley, 1992). Despite these efforts, the results have proven controversial regarding the existence of the Kuleshov effect (Barratt et al., 2016; Prince & Hensley, 1992).

Initially, Prince and Hensley (Prince & Hensley, 1992) attempted to replicate the Kuleshov Experiment paradigm. In Prince’s study, each sequence started with a fade-in on an actor’s neutral face, followed by a cut to an object (soup, coffin, or child), and then a cut back to the actor’s neutral face, concluding with a fade-out. Prince tasked the participants with discerning whether the actor’s neutral face conveyed emotion. However, the majority of the participants perceived no emotion in the actor, and Prince’s study did not confirm the existence of the Kuleshov effect.

In a subsequent study, Ildirar and Ewing (Ildirar & Ewing, 2018) replicated the Kuleshov effect experiment, categorizing participants into two groups: first-time viewers and experienced viewers. Ildirar edited emotional facial clips and emotional contextual clips together to investigate whether viewers constructed spatiotemporal links between the two clips, recognizing that the two clips conveyed a unified narrative. The study revealed that experienced viewers were able to construct these links, while first-time viewers could not. This finding provided confirmation of the existence of the Kuleshov effect.

While several earlier studies predominantly explored the Kuleshov effect from a film-centric perspective (Ekman, 1971; Ildirar & Ewing, 2018; Prince & Hensley, 1992), more recently investigations have employed the Kuleshov paradigm to scrutinize the contextual framing impact on facial emotion recognition from psychological perspective. Barratt et al. (Barratt et al., 2016) conducted a study that investigated the influence of emotional contexts on facial stimuli using the Kuleshov paradigm. They refined the experimental materials by incorporating neutral faces from the Karolinska Directed Emotional Faces (KDEF) picture set as facial stimuli and integrating online static or dynamic videos as varied emotional stimuli. Employing the Kuleshov sequence, participants were tasked with evaluating the valence and arousal of the actor. Barratt discovered that participants’ judgments of facial expressions were indeed influenced by the emotional stimuli, thus affirming the existence of the Kuleshov effect. Subsequently, Calbi et al. (Calbi et al., 2017) enhanced the ecological validity of Barratt’s experiment by exclusively utilizing dynamic images for reverse shots and introducing inter-trial intervals. Participants initially viewed a Kuleshov sequence, and then they were asked to rate the valence of the neutral face. The results revealed that the emotional context affected the emotional judgment of neutral faces, resulting in a context-dependent bias of emotion perception. In a similar vein, Mullennix et al. (Mullennix et al., 2019) rigorously manipulated the Kuleshov effect paradigm, utilizing static facial stimuli and varying presentation times to investigate the effect of visual context on the interpretation of facial expressions. After exposure to an emotional context, participants viewed a neutral face, and the results indicated that more faces were labeled as “happy” after viewing a pleasant context and more faces were labeled “sad” or “fearful” after viewing an unpleasant context. In summary, these studies collectively reveal that the assessment of neutral faces’ emotional expressions is influenced by the preceding emotional context when using static faces and online emotional videos. This suggests that the Kuleshov effect can be detected with conventional psychological experiment materials.

However, when revisiting the Kuleshov effect, the significant disparity between static images and dynamic films, in accordance with the definition of film employing moving images to create continuous life experiences (Fiorelli, 2016), cannot be disregarded. Specifically, although the use of zoom-ins with static neutral faces from the KDEF picture set approximates reality (Calbi et al., 2017), it fails to capture the nuanced facial movements inherent in actual film scenes (Ambadar et al., 2005; Tan, 2018), resulting in the omission of spatiotemporal information from dynamic faces (Krumhuber et al., 2023). Moreover, in authentic films, the continuity of backgrounds between scenes establishes a cohesive spatial context that enables viewers to perceive these scenes as unfolding within the same spatial realm (Carroll & Carroll, 1993; T. J. Smith, 2012). The replication studies predominantly rely on isolated facial expressions and emotional scenes (Calbi et al., 2017; Ildirar & Ewing, 2018; Mullennix et al., 2019), overlooking the alignment between facial backgrounds and emotional scenes present in real-world film scenarios. This departure from authentic film scenarios may potentially compromise the robustness of the validation of the Kuleshov effect. Therefore, there is a compelling need to reexamine the Kuleshov effect using high ecological film materials.

In addition to revisiting the Kuleshov effect through behavioral experiments, employing cutting-edge neuroimaging techniques can bolster its confirmation by revealing its neural correlates. This approach aligns with the neurocinematic exploration of films (Hasson et al., 2008). To our knowledge, only two studies have attempted to explore the neural correlates of the Kuleshov effect. Mobbs et al. (Mobbs et al., 2006) conducted a functional magnetic resonance imaging (fMRI) experiment that unveiled specific mechanisms involved in the Kuleshov effect, particularly regarding how emotional scenes impact neutral facial expressions. Through the juxtaposition of various emotional scenes with neutral facial expressions in a scene-face sequence, the study identified activation in distinct brain regions, including bilateral temporal pole (TP), anterior cingulate cortex (ACC), amygdala, and bilateral superior temporal sulcus (STS). This research shed light on the neural correlates of the Kuleshov effect. Similarly, Calbi et al. (Calbi et al., 2019) employed EEG and discovered that viewers rated faces as more arousing when preceded by a fearful context. The study pinpointed a high-amplitude late positive potential (LPP) in the EEG data when faces were preceded by emotional contexts, indicating a cognitive process of expectation attribution influencing how facial expressions were interpreted. This was evidenced by the activation of LPP when evaluating incongruent sequences of stimuli. However, these studies have not addressed the neural correlates of the Kuleshov effect within a face-scene-face sequence using authentic films. This gap in research makes it challenging to fully understand how the Kuleshov effect is generated during the process of film watching.

In this study, our objective is to investigate the presence of the Kuleshov effect using authentic films. In Experiment 1, we validated the existence of the Kuleshov effect with authentic films, employing film cameras to capture actors’ neutral facial expressions and emotional scenes. These elements were combined into a face-scene-face sequence, ensuring consistency in the background between faces and emotional scenes in authentic film clips. Participants were then tasked with rating the emotions portrayed by the actors. Our hypothesis posited that the judgment of emotion in neutral faces would be influenced by the emotional scene (fearful, neutral, happy). In Experiment 2, we sought to explore the neural correlates of the Kuleshov effect to further validate its existence. Employing identical authentic films, we refined the Kuleshov sequence by incorporating jetters into the procedure to accommodate fMRI data acquisition. Our hypothesis postulated that the inclusion of jetters would not affect the observation of context-dependent bias. Furthermore, we aimed to observe the neural correlates of the Kuleshov effect by contrasting activation between faces. Subsequently, we examined the context-dependent bias of Kuleshov effect, investigating how brain activity elicited by faces was altered by emotional scenes.

## 2. Methods

The current study adopted the Kuleshov experimental paradigm (Calbi et al., 2017), which includes a neutral face, an emotional scene, and a neutral face presented in a face-scene-face sequence. The aim was to observe the context-dependent bias in judging neutral faces affected by accompanying emotional scenes (fearful, neutral, happy) within authentic films. In Experiment 1, we captured authentic face and scene materials and combined them in film sequences. Subsequently, these film sequences were employed in a behavioral experiment to explore the existence of the Kuleshov effect. In Experiment 2, the same films were utilized in an fMRI study to investigate the neural correlates of the Kuleshov effect.

One hundred and two healthy volunteers participated in the experiments, and all provided informed consent, which was approved by the ethics committee of the State Key Laboratory of Cognitive Neuroscience and Learning at Beijing Normal University. Participants received monetary compensation, and the experimental procedures were ethically sanctioned by the aforementioned committee.

### 2.1. Experiment 1

#### 2.1.1. Participants

Seventy-one healthy volunteers (35 females) with either normal vision or vision corrected to normal were recruited from Beijing Normal University. Twelve participants (6 females) participated in the materials rating experiment, while fifty-nine participants (29 females) were involved in the Kuleshov experiment. Eligible participants, as confirmed through a questionnaire, reported the absence of panic disorder. Individuals majoring in film studies were deliberately excluded from participation.

#### 2.1.2. Stimuli

To create authentic films for the Kuleshov effect, we meticulously designed the camera positions to adhere to the face-scene-face sequence utilizing the shot-reverse-shot structure (Figure 1A). Following the established Kuleshov experiment paradigm, which includes a neutral face, an emotional scene, a repeated neutral face, we captured one set of face clips and one set of emotional scene clips. All clips were then converted to grayscale, and sound was removed to eliminate the potential impact of color and sound, in line with prior studies (Barratt et al., 2016; Calbi et al., 2017). Subsequently, we seamlessly combined them in a face-scene-face sequence utilizing shot-reverse-shot structure, presenting the actor’s face in the “shot” and the corresponding emotional scene in the “reverse shot” to reflect what the actor is supposedly seeing.

**Figure 1.**
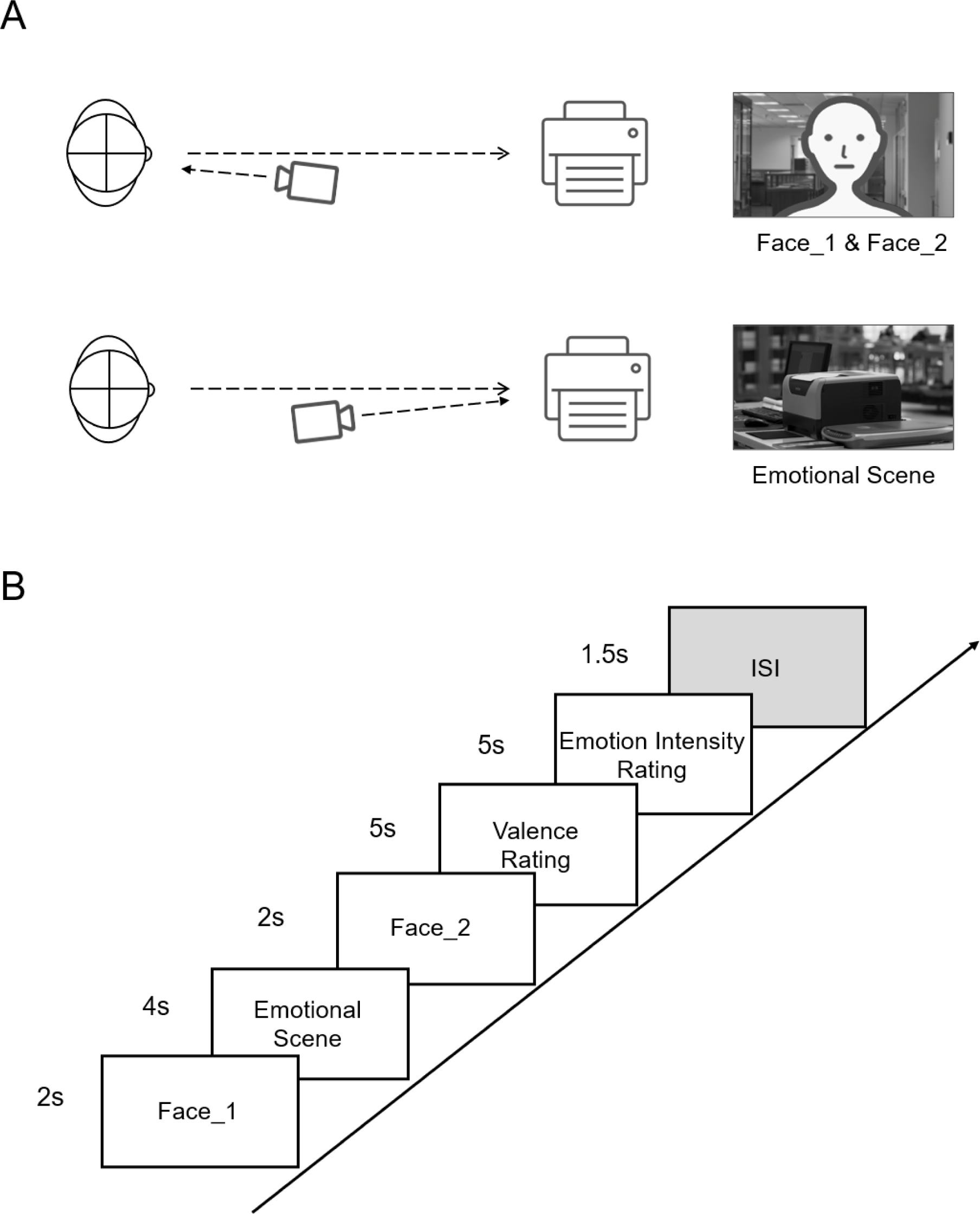
Experimental Materials and Procedure. (A) In this figure, we present the depiction of camera positions in the Kuleshov sequence. In the case of neutral Face_1 and Face_2, the actor, depicted with a stick figure to preserve privacy, was specifically directed to gaze at the object without expressing any emotion. Conversely, for the emotional scene, serving as a reverse shot, the camera was strategically positioned near the 180-degree axis, directed towards the object. (B) Each trial started with a 2-second clip featuring a neutral face, followed by a 4-second clip depicting an emotional scene, and concluded with another 2-second clip of the same neutral face. After viewing the film, participants were instructed to assess the valence and emotional intensity of the neutral face. The trial concluded with a 1.5-second ISI.

##### Neutral Faces

For the face clips, we employed a cinecamera, lighting equipment, and a blue screen for the film shoot. Actors were seated in front of the blue screen and were instructed to maintain a neutral facial expression while looking at a focal point adjacent to the camera. Throughout the recording, actors kept their facial muscles relaxed, minimizing movements to blinking only. We recorded 116 face clips from various actors, each cut into a 2-second clip. Subsequently, we carefully selected 15 male and 15 female actors based on their neutral expressions and ensured similarity in appearance. This selection was achieved through an internal group rating conducted by the authors. The 30 neutral faces were randomly allocated into three groups to construct the emotional conditions in the Kuleshov experiment. To assess the emotional expression of these neutral faces, a facial rating experiment was conducted, with detailed information shown in the Supplementary Materials. Briefly, twelve participants rated the valence of the actors’ neutral face. The averaged valence approached zero, at 0.02 ± 0.13. Following this, the faces were randomly assigned to three groups: fearful, neutral, and happy. No significant differences were observed among the three groups (*F*_1,2_ = 0.58, *p* = 0.569).

##### Emotional scenes

To capture emotional scenes, we collaborated with a professional film director and a dedicated crew. Initially, we carefully designed these scenes before shooting, ensuring their compatibility with the shot-reverse-shot structure while maintaining a consistent 180-degree axis orientation (Figure 1A). Three distinct categories of emotional scenes were conceptualized, each encompassing 10 unique contexts (Figure S1). These scenarios spanned various genres, including horror, documentary and comedy. Fearful scenes depicted chilling scenarios such as murder, ghosts, and peeping. Neutral scenes featured commonplace settings like a bus stop, a printer, and a traffic light. Happy scenes showcased joyful images of food, an attracted girl, and humorous expressions. The production quality for each emotional scene was meticulously maintained at a cinematic level, involving coordination with the film director, lighting setup, acting, and cinematography. After capturing the raw footage, we implemented color correction to impart a cinematic feel to the videos. Following this, the videos were edited into 4-second clips to serve as emotional scene materials in the Kuleshov experiment. The clip length aligns with the Average Shot Length (ASL) found in mainstream Hollywood films, typically ranging between 3 and 4 seconds (Calbi et al., 2017; Cutting et al., 2011; Salt, 1974). To assess the emotional impact of these scenes, a scene rating experiment was conducted, involving the same twelve participants who took part in the facial rating experiment (See Supplementary Materials). The results indicated negative valence for fearful scenes (-2.19 ± 0.24), positive valence for happy scenes (1.98 ± 0.17), and neutral valence for neutral scenes (0.48 ± 0.08), with significant differences observed among the three groups (*F*_1,2_ = 146.88, *p* < 0.001).

##### Final stimuli

For each trial in the Kuleshov experiment, we selected a clip from the neutral facial materials and a clip from the emotional scenes, combining them in the sequence of face-scene-face, ensuring each face and emotional scene was utilized only once. This resulted in a total of 30 combined films (Figure 1A). To maintain gender balance of neutral faces, each emotional scene included 5 male and 5 female actors. To ensure sexual attraction between genders, we employed cross-gender matching, pairing male faces with female emotional scenes and vice versa. To creating a more authentic film scenario, it was crucial to ensure that the backgrounds among different video clips were similar. This background matching not only enhanced the ecological design of the Kuleshov validation experiment but also addressed a previously overlooked aspect in similar studies (Calbi et al., 2017; Ildirar & Ewing, 2018). In the matching process, blue screen image matting was utilized to extract the actors, and subsequently, the backgrounds were substituted with videos or images that closely resembled emotional scenes. Notably, most videos and images were captured simultaneously with the emotional scenes. Adjustments were made to the color temperature, brightness, and contrast of the facial videos to harmonize them with the emotional scenes.

#### 2.1.3. Procedure

In each trial, participants initiated the session by viewing a 2-second clip featuring a neutral face, succeeded by a 4-second clip of an emotional scene, and concluding with another 2-second clip of the same neutral face (Figure 1B). Following the 8-second film presentation, participants were tasked with rating the actor’s performance emotions using two distinct scales. The first scale involved rating the valence of the neutral face on a scale from -1 to 1, where -1 denoted a fearful emotion, 0 represented a neutral emotion, and 1 indicated a happy emotion. On the second scale, participants rated the emotional intensity of the neutral face on a scale from 1 to 5, with 1 signifying low emotional intensity and 5 representing high emotional intensity. Each rating had a 5-second response time. Subsequently, a 1.5-second inter-stimuli interval (ISI) concluded the trial, and participants seamlessly progressed to the subsequent trial, completing a total of 30 trials (Figure 1A).

Stimuli delivery and response recording were controlled using PsychoPy 3.2 software (https://www.psychopy.org/), administered on a 14-inch laptop. To minimize potential influences on the Kuleshov effect, participants were queried with two questions upon completing the experiment: (1) “Have you heard of the Kuleshov effect?” and (2) “How many movies do you watch per week on average?” Additionally, to maintain balanced emotional conditions, half of the participants followed the sequence ’fearful – neutral – happy,’ while the other half followed ’happy – neutral – fearful.’ Gender distribution was also balanced among the subgroups.

#### 2.1.4. Data Analysis

The Kuleshov effect, known for influencing viewers’ interpretations of actors’ facial expressions based on emotional scenes, was examined in this study. Our hypothesis posits that the existence of the Kuleshov effect would be evident if emotional scenes influence the rating of emotions in neutral faces, leading to a valence score different from 0. More precisely, we anticipate that fearful scenes will prompt a negative emotional rating for neutral faces (valence score tending to -1), while happy scenes will evoke a positive emotion (valence score tending to 1). To test this, we computed the average valence for all participants under each type of emotional condition, serving as an indicator of the degree of the Kuleshov effect. Subsequently, a one-way analysis of variance (ANOVA) was conducted to identify statistical differences in valence among the three emotional conditions. *Post hoc* comparisons among means were performed using LSD’s test. A similar analytical approach was employed for emotional intensity.

### 2.2 Experiment 2

#### 2.2.1. Participants

Thirty-one healthy volunteers (16 females, age 21.94 ± 2.99 years) with normal or corrected-to-normal vision were recruited from Beijing Normal University. Participants were screened for the absence of panic disorder through a questionnaire, and individuals majoring in film studies were excluded.

#### 2.2.2. Procedure

In this experiment, authentic films from Experiment 1 were employed. To distinguish blood-oxygen-level-dependent (BOLD) activity between neutral faces and emotional scenes, the Kuleshov sequence was modified by introducing randomized jitters between neutral faces and emotional scenes (Figure 2).

**Figure 2.**
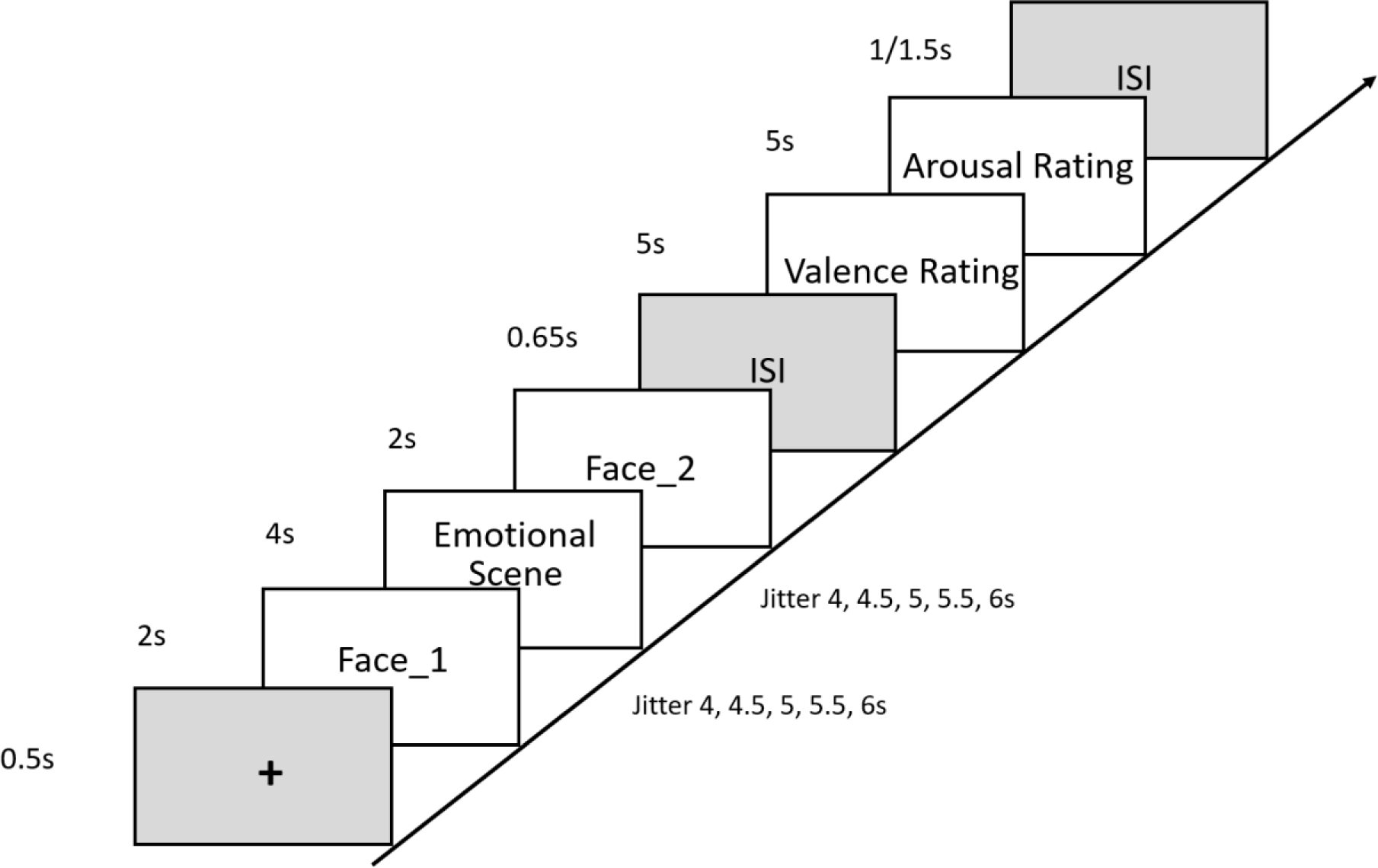
Experimental Procedure in Experiment 2. Each trial initiated with a 0.5-second presentation of a crosshair sign, succeeded by a 2-second presentation of a neutral face. This was followed by a variable jitter lasting 4 to 6 seconds, which in turn was followed by a 4-second presentation of an emotional scene. Another jitter lasting 4 to 6 seconds followed, and then, a 2-second presentation of a neutral face. Following the film presentation, a 0.65-second ISI ensued, after which participants were required to rate valence and arousal. Each rating had a 5-second response time. The trial concluded with a 1 to 1.5-second ISI.

Before the experiment, participants completed two practice trials to familiarize themselves with the procedure, using films different from those in the formal experiment. Subsequently, participants underwent T1-weighted image scans and then completed the Kuleshov sequence during fMRI scans.

In each trial, participants first viewed 10 seconds of experiment instructions before the first trial. Each trial initiated with a 0.5-second crosshair sign, followed by 2 seconds of neutral face presentation. A random jitter lasting 4 to 6 seconds ensued, succeeded by 4 seconds of emotional scene presentation, another 4 to 6 seconds of a jitter, and then, 2 seconds of neutral face presentation. Following the film presentation, there was a 0.65-second ISI. Participants then rated valence and arousal of the neutral face. Each rating had a 5-second response time. The valence scale ranged from -4 (negativity) to 4 (positivity), and the arousal scale ranged from 1 (low arousal) to 9 (high arousal). Subsequently, an ISI lasting 1 to 1.5 seconds concluded the trial. completing one trial, participants proceeded to the next, resulting in a total of 30 trials.

Stimuli delivery and response recording were controlled using PsychoPy 3.2 software. To minimize potential influences on the Kuleshov effect, participants were queried at the conclusion of the experiment: “Have you heard of the Kuleshov effect?” Additionally, to balance the order of emotional conditions, the presentation order of different films was randomized across participants.

#### 2.2.3. Data Acquisition

In this experiment, the Siemens 3T Prisma MRI scanner was utilized for scanning. Magnetization Prepared Rapid Acquisition Gradient-echo (MPRAGE) imaging was employed to scan the 31 participants and acquire their three-dimensional T1-weighted data. The imaging parameters were as follows: repetition time (TR) / echo time (TE) / inversion time (TI) = 2530 ms / 2.27 ms / 1100 ms; flip angle (FA) = 7°; field of view (FOV) = 256 × 256 mm^2^; number of slices = 208; slice thickness = 1 mm; voxel size = 1 × 1 × 1 mm^3^.

Task-related fMRI scans were performed using a T2-weighted echo planar imaging sequence (EPI/separate). The imaging parameters were as follows: TR / TE = 2000 ms / 34 ms; FA = 70°; FOV = 200 × 200 mm^2^; matrix size = 64 × 64 mm^2^; number of slices = 72; slice thickness = 2 mm; slice gap = 0 mm; voxel size = 2 × 2 × 2 mm^3^. The average acquisition time for structural and functional image data was 25 minutes and 12 seconds.

#### 2.2.4. Data Analysis

##### Behavioral data

In analyzing the behavioral data, the primary focus of the current study was to explore the impact of introducing jitters between neutral faces and emotional scenes on observing the Kuleshov effect. This modification in the Kuleshov sequence aligns with film editing techniques, such as the use of a black screen in thriller films (Demme, 1991). The analytical approach for behavioral data mirrored that of Experiment 1. Average valence and arousal scores were calculated for all participants across each type of emotional condition. Statistical differences among the three emotional conditions were assessed using repeated-measures analysis of variance (ANOVA), and *post hoc* comparisons among means were conducted using LSD’s test.

##### Preprocessing of fMRI data

In this experiment, we conducted preprocessing and analysis of the brain imaging data from the 31 participants using MRIcron software (https://www.nitrc.org/projects/mricron/) and SPM12 software (https://www.nitrc.org/projects/spm/). The preprocessing steps included the following: 1) Conversion of scanned DICOM format data to NIfTI format. 2) Time-slice correction to rectify time differences between different slices. 3) Field map correction to address geometric distortions caused by magnetic field inhomogeneity. Two different echo times (TE) of 4.92 milliseconds and 7.38 milliseconds were used, and distortion correction was performed using phase and magnitude images. 4) Head motion correction and distortion correction, with a motion correction setting of a quality parameter at 0.9 and a 4mm separation, Gaussian kernel parameters (FWHM 5mm × 5mm × 5mm). 5) Registration of structural and functional images, utilizing normalized mutual information (nmi) as the cost function and setting spatial separation parameters and Gaussian kernel parameters (FWHM 7mm × 7mm × 7mm). 6) Segmentation of T1 images into different tissue types such as scalp, skull, cerebrospinal fluid, gray matter, and white matter. 7) Spatial normalization, aligning functional images to the MNI standard template and resampling to 2mm × 2mm × 2mm. 8) Spatial smoothing with Gaussian kernel parameters (FWHM 4mm × 4mm × 4mm) to enhance signal-to-noise ratio for group-level analysis.

##### Analysis of fMRI data

In the fMRI data analysis, our investigation into the neural correlates of the Kuleshov effect comprised three primary objectives. Firstly, we aimed to identify neural correlates associated with the creation of new meaning on the neutral face within the Kuleshov sequence. To achieve this, we contrasted the neural activity evoked by Face_2 and Face_1 in each condition, assuming the existence of the Kuleshov effect. Secondly, we sought to uncover the neural correlates of the Kuleshov effect bias, which leads to a context-dependent bias in emotional perception. This was achieved by comparing the neural activity elicited by Face_2 in the fearful or happy condition with that in the neutral condition. Thirdly, we explored the interaction between the neutral face processing and emotional conditions by examining the (Face_2 – Face_1) contrast in the fearful or happy condition against the (Face_2 – Face_1) contrast in the neutral condition. This allowed us to detect unique brain regions contributing to negative or positive emotional perception bias, providing further evidence for the existence of the Kuleshov effect.

To accomplish these objectives, we conducted first-level SPM analyses on each participant using a general linear model to obtain contrast activation. The first-level SPM analysis involved averaging activation differences across 10 trials of the same emotional condition. Subsequently, these contrast images were entered into second-level SPM analyses (one-sample T-tests) to evaluate the main effect of each contrast. The results of the second-level analysis were obtained in the form of spm_T maps. Subsequently, further statistical analysis was conducted using xjview software (https://www.alivelearn.net/xjview/). This involved selecting positively activated brain regions, establishing a cluster threshold of 5, applying false discovery rate (FDR) correction for multiple comparisons, and setting the significance level at *p* < 0.05. Following these steps, we reported the locations of brain regions with FDR multiple comparison correction.

## 3. Results

### 3.1. Experiment 1

In Experiment 1, our primary goal was to ascertain the existence of the Kuleshov effect, which manifests as the perception of emotions from neutral faces influenced by the inserted emotional scenes. During the material rating experiment, neutral faces received an average valence rating close to zero (0.02 ± 0.13), indicating a neutral baseline.

Upon incorporating these neutral faces into the Kuleshov sequence, the emotional perception of these neutral faces was significantly affected by the accompanying emotional scene. Specifically, in the fearful condition, exposure to fearful scenes led to a negative valence rating for neutral faces (-0.45 ± 0.04), indicating a perceived negative emotion. Conversely, in the happy condition, where scenes evoked positive emotions, a positive valence rating for neutral faces was observed (0.27 ± 0.04). Importantly, in the emotionless condition, neutral scenes did not induce any emotional perception, reflected in a neutral valence rating for neutral faces (0.07 ± 0.03), as illustrated in Figure 3A. The results affirm that viewers attribute emotions to neutral faces.

**Figure 3.**
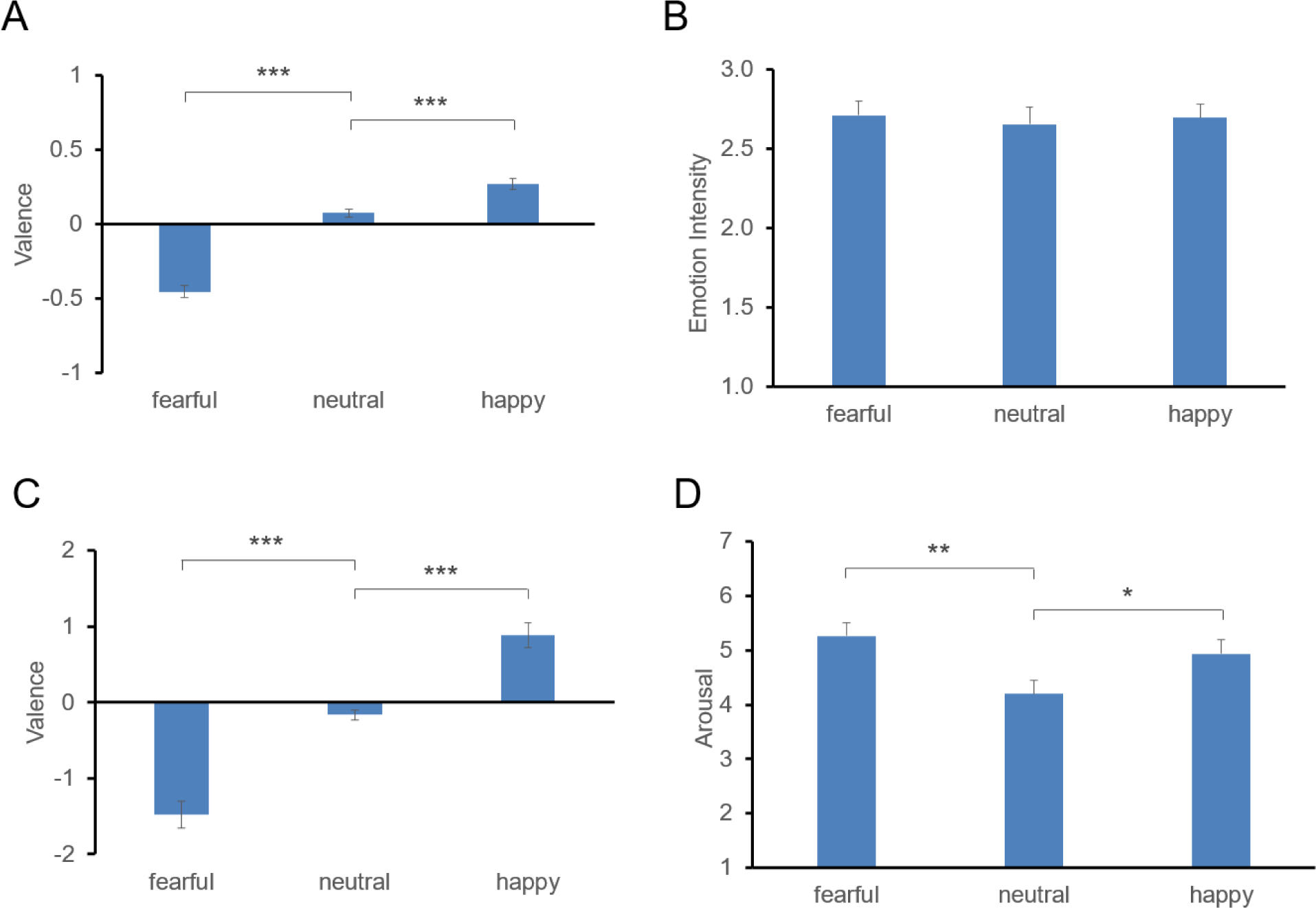
Behavioral results. (A) Averaged valence scores from 59 participants in Experiment 1. A significant main effect of emotional conditions is observed (*F*1,2 = 116.05, *p* < 0.001). (B) Averaged emotional intensity scores from 59 participants in Experiment 1. No significant difference is found among emotional conditions. (C) Averaged valence scores from 31 participants in Experiment 2. A significant main effect of emotional conditions is evident (*F*1,2 = 70.48, *p* < 0.001). (D) Averaged arousal scores from 31 participants in Experiment 2. A significant main effect of emotional conditions is detected (*F*1,2 = 4.91, *p* = 0.009). (\**p* < 0.05, \*\**p* < 0.01, \*\*\**p* < 0.001)

To assess whether the Kuleshov effect introduced a context-dependent bias in emotional perception of neutral faces, we conducted an ANOVA on the three emotional conditions (fearful, neutral, happy), revealing a significant main effect (*F*_1,2_ = 116.05, *p* < 0.001). *Post hoc* tests confirmed significant differences in valence within each condition (*p* < 0.001), supporting the existence of the Kuleshov effect.

Despite emotional scenes influencing valence ratings, no parallel bias emerged in emotional intensity ratings. The main effect of emotional scenes on emotional intensity was not significant (*F*_1,2_ = 0.09, *p* = 0.913, shown in Figure 3B), indicating that participants perceived similar emotional intensity in actors’ performances across different emotional scenes. Lastly, prior knowledge of the Kuleshov effect or viewing frequency does not impact the observation of the Kuleshov effect (Figure S2 and Figure S3).

### 3.2. Experiment 2

#### 3.2.1. Behavioral data

In this analysis, we sought to validate the existence of the Kuleshov effect using a modified paradigm that introduced jitters between scenes and faces. Our findings indicate that the inclusion of jitters did not diminish the Kuleshov effect. Fearful scenes elicited a negative valence rating on neutral faces (-1.48 ± 0.17), while happy scenes generated a positive valence rating (0.89 ± 0.16). However, neutral scenes did not induce an emotional bias in valence ratings for neutral faces (-0.16 ± 0.07), as illustrated in Figure 3C. Subsequently, an ANOVA confirmed a significant main effect of emotional scenes (*F*_1,2_ = 70.48, *p* < 0.001), with post hoc tests indicating significant differences in valence within each condition (*p* < 0.001), reaffirming the existence of the Kuleshov effect in the modified sequence.

Furthermore, an analysis of emotional arousal revealed that fearful scenes evoked the highest arousal levels (5.27 ± 0.24), followed by happy scenes (4.94 ± 0.25), while neutral scenes registered the lowest arousal levels (4.20 ± 0.25), as depicted in Figure 3D. ANOVA results demonstrated a significant main effect of emotional scenes on arousal (*F*_1,2_ = 4.91, *p* = 0.009). *Post hoc* tests indicated significant arousal differences between fearful and neutral conditions (*p* = 0.003) and between happy and neutral conditions (*p* = 0.037). These findings suggest that emotional scenes triggered more robust emotional responses compared to neutral scenes. Importantly, none of the participants reported prior knowledge of the Kuleshov effect.

#### 3.2.2. fMRI data

##### Direct comparison of Face_2 and Face_1 in each condition

In the Kuleshov sequence, viewers first see a neutral face, followed by an emotional scene, and then a similar neutral face. Subsequently, viewers attribute emotion to the neutral face (Figure 3), suggesting a perceived new meaning for the second neutral face. Detecting neural correlates of this phenomenon is crucial for affirming the existence of the Kuleshov effect.

To uncover the neural correlates associated with the new meaning attributed to the second face, our fMRI analysis compared brain activity between Face_2 and Face_1. We hypothesized that Face_2, when contrasted with Face_1, would display heightened activation across the entire brain, with distinct regions being activated under different emotional conditions.

In summary, we observed widespread activation across the entire brain in each emotional condition, but the specific activated regions varied (Figure 4). In the fearful condition, subtracting Face_1 from Face_2 revealed 46 clusters, including bilateral activation in the hippocampus, insula, precentral gyrus, postcentral gyrus, ACC, posterior cingulate cortex (PCC), orbitofrontal cortex (OFC), inferior parietal lobe (IPL), and cuneus (for additional AAL atlas labels, refer to Table S1 and Figure 4B). These regions are implicated in emotional memory and attentional control, crucial for the Kuleshov effect in fearful conditions. Similarly, in the happy condition, subtracting Face_1 from Face_2 identified 51 clusters with bilateral activation in the insula, middle frontal gyrus (MFG), ACC, midcingulate cortex (MCC), PCC, IPL, and precuneus (for additional AAL atlas labels, refer to Table S2 and Figure 4D). These activated regions underscore the role of emotional memory and attentional control in the Kuleshov effect during happy conditions. In the neutral condition, subtracting Face_1 from Face_2 revealed 55 clusters with bilateral activation in the supplementary motor area (SMA), postcentral gyrus, precentral gyrus, IPL, MFG, ACC and PCC (for additional AAL atlas labels, refer to Table S3 and Figure 4C). These activated regions suggest the involvement of motor control and sensory information processing in the Kuleshov effect during neutral conditions.

**Figure 4.**
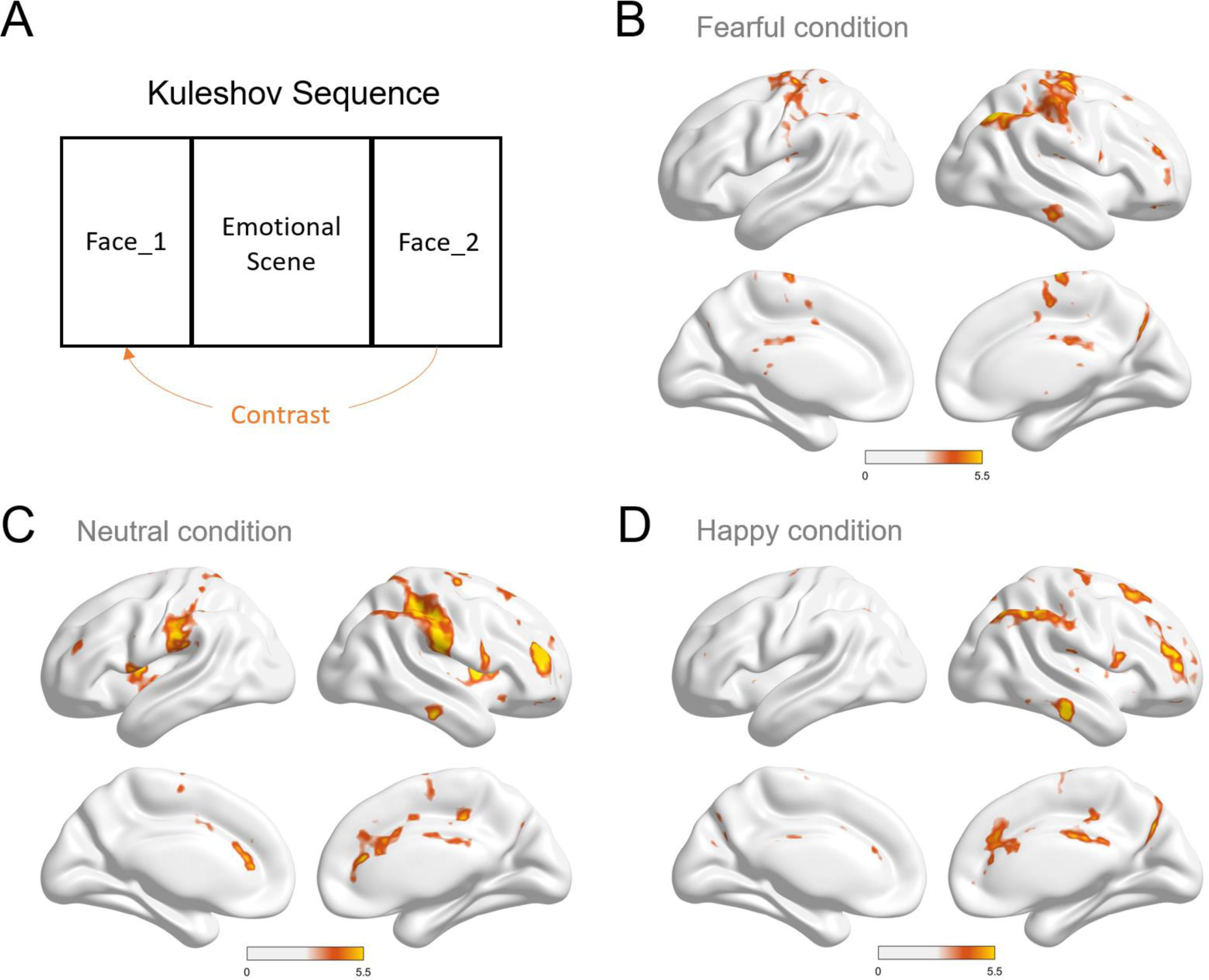
Comparison of brain activity between Face_2 and Face_1 in each emotional condition. Brain activity was obtained by subtracting Face_1 from Face_2 (A), and the contrasts were conducted for the fearful condition (B), neutral condition (C), and happy condition (D) (*p* < 0.05, FDR-corrected, cluster size > 5 voxels).

##### Direct comparison of Face_2 between fearful or happy condition and neutral condition

Consistent with behavioral findings, the Kuleshov effect introduced a discernible context-dependent bias in emotional perception, altering the interpretation of neutral faces based on preceding emotional scenes (Figure 3). Our subsequent analysis aimed to probe the neural correlates of this Kuleshov effect bias, focusing on how neutral faces exhibit distinct patterns of brain activation when preceded by either fearful or happy scenes (Mobbs et al., 2006).

In the fearful condition compared to the neutral condition, the contrast between Face_2 revealed significant activation in 17 clusters, involving regions such as the bilateral cerebellum, fusiform gyrus (FG), parahippocampal gyrus (PHC), angular gyrus (AG), PCC, cuneus, precuneus, precentral gyrus, and postcentral gyrus (for additional AAL atlas labels, Table 1 and Figure 5A). These regions are well-established in facial and emotional processing, supporting the notion that the Kuleshov effect reflects a contextual modulation of face perception.

**Table 1.** fMRI Results: Face_2 in fearful or happy condition minus Face_2 in neutral condition.

| Brain Region | AAL Atlas Labels | Peak Voxel Coordinate (MNI) | Cluster Size (KE) | T-score |
| --- | --- | --- | --- | --- |
| ***Fearful > Neutral*** (FDR-corrected cluster threshold, $p < 0.05$) | | | | |
| Right Cerebellum | Cerebellum_9_L | -2, -52, -40 | 6 | 5.259 |
| Left Cerebellum | Cerebellum_Crus2_L<br>Cerebellum_Crus1_L | -40, -72, -36 | 8 | 4.878 |
| Left Fusiform Gyrus | Fusiform_L | -36, -46, -20 | 7 | 5.232 |
| Left Middle Temporal Gyrus | Temporal_Mid_L | -52, -12, -18 | 9 | 6.367 |
| Right ParaHippocampal Gyrus | ParaHippocampal_R<br>Fusiform_R | 34, -32, -14 | 27 | 7.138 |
| Right Lingual Gyrus | Lingual_R<br>Occipital_Inf_R | 26, -88, -14 | 6 | 4.729 |
| Right VLPFC | Frontal_Inf_Orb_2_R | 52, 40, -6 | 11 | 6.107 |
| Left Precuneus | Precuneus_L<br>Calcarine_L | -10, -60, 16 | 5 | 5.144 |
| Right Precuneus | Precuneus_R<br>Cuneus_R<br>Calcarine_R | 10, -62, 24 | 21 | 5.190 |
| Left Posterior Cingulate Cortex/Precuneus | Precuneus_L<br>Precuneus_R<br>Cingulate_Post_L | 0, -56, 20 | 22 | 5.598 |
| Left Angular Gyrus | Angular_L<br>Occipital_Mid_L<br>Temporal_Mid_L | -42, -70, 28 | 39 | 5.180 |
| Left Lateral Occipital Cortex | Occipital_Mid_L | -32, -84, 34 | 8 | 5.168 |
| Right Angular Gyrus | Angular_R | 38, -68, 42 | 11 | 4.607 |
| Precentral Gyrus | Precentral_R<br>Postcentral_R | 30, -22, 52 | 28 | 5.461 |
| Right Superior Frontal Sulcus | Frontal_Sup_2_R<br>Frontal_Mid_2_R | 30, 32, 54 | 16 | 5.802 |
| Left Precentral Gyrus | Precentral_L | -30, -24, 64 | 21 | 5.304 |
| Right Postcentral Gyrus | Precentral_L<br>Postcentral_L | -38, -26, 62 | 5 | 4.969 |
| ***Happy > Neutral (FDR-corrected cluster threshold, $p < 0.05$)*** |  |  |  |  |
| Right Precuneus/Cuneus/CAL | Precuneus_R<br>Cuneus_R<br>Calcarine_R | 18, -56, 20 | 31 | 6.314 |

**Figure 5.**
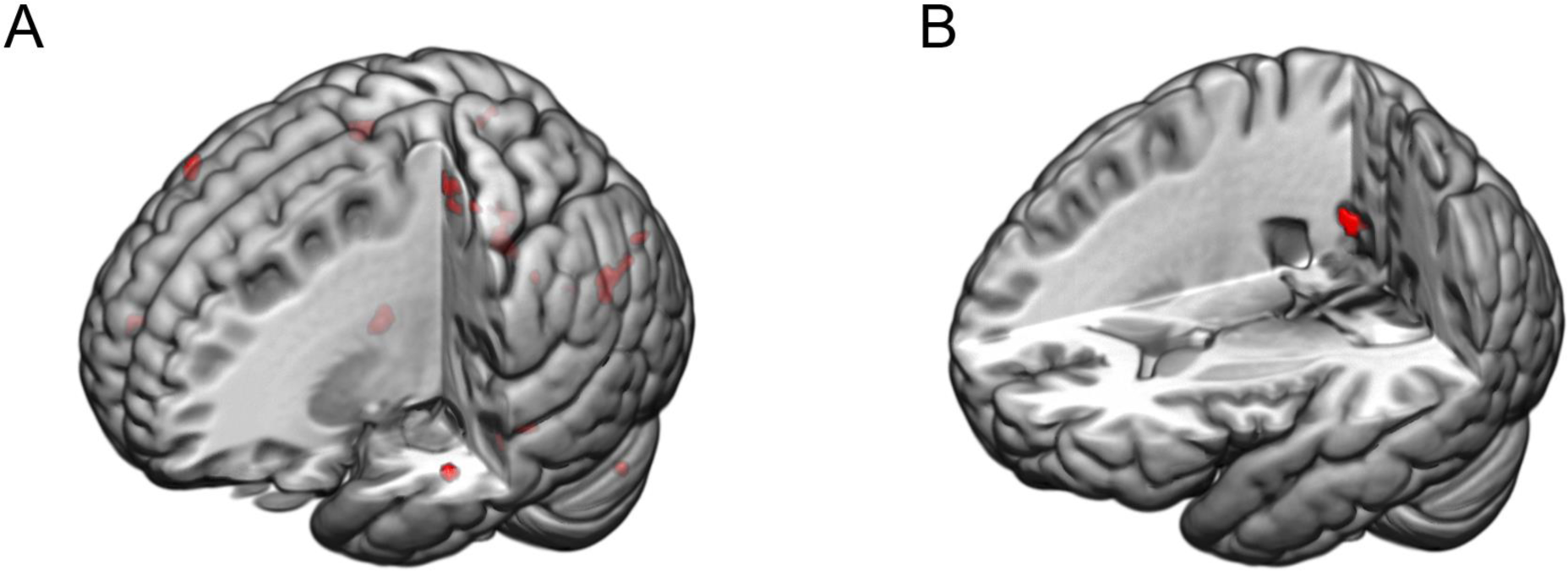
Comparison of brain activity for Face_2 between emotional and neutral conditions. (A) Depicts brain regions illustrating the primary effect of Face_2 in the fearful condition compared to Face_2 in the neutral condition. (B) Illustrates brain regions showing the primary effect of Face_2 in the happy condition compared to Face_2 in the neutral condition (*p* < 0.05, FDR-corrected, cluster size > 5 voxels).

Conversely, when contrasting Face_2 in the happy condition with those in the neutral condition, only one cluster of significant activation emerged, encompassing the right cuneus, right precuneus, and right calcarine fissure and surrounding cortex (CAL) (Table 1 and Figure 5B). These regions are linked to visual processing, suggesting that the influence of happy scenes on neutral faces is comparatively weaker than that of fearful scenes.

##### Interaction between neutral face processing and emotional conditions

Viewers perceive negative emotions from neutral faces in fearful conditions and positive emotions in happy conditions, as illustrated in Figure 3B, suggest an interaction between neutral facial processing and emotional conditions. Viewers perceive negative emotions from neutral faces in fearful conditions and positive emotions in happy conditions. This dynamic interaction adds a crucial layer to our understanding of the Kuleshov effect and its intricate relationship with context-dependent biases.

Examining the (Face_2 – Face_1) contrast in the fearful condition against the neutral condition revealed activation in 10 clusters, including the right insula, bilateral precentral gyrus, bilateral postcentral gyrus, right supplementary motor area (SMA), and bilateral paracentral lobule (Table 2 and Figure 6A). These specific brain regions provide insights into the neural substrates of the Kuleshov effect in fearful scenes.

**Table 2.** fMRI Results: interaction between neutral face processing and emotion conditions.

| Brain Region | AAL Atlas Labels | Peak Voxel Coordinate (MNI) | Cluster Size (KE) | T-score |
| --- | --- | --- | --- | --- |
| *Fearful [Face_2 – Face_1] > Neutral [Face_2 – Face_1] (FDR-corrected cluster threshold, $p < 0.05$)* |  |  |  |  |
| Right Insula/Rolandic Operculum | Insula_R<br>Rolandic_Oper_R | 36, -18, 18 | 6 | 4.813 |
| Left Precentral Gyrus | Precentral_L | -24, -24, 54 | 44 | 6.032 |
| Right Precentral Gyrus | Precentral_R | 28, -24, 54 | 34 | 6.200 |
|  | Postcentral_R |  |  |  |
| Left Postcentral Gyrus/Precentral Gyrus | Postcentral_L<br>Precentral_L | -38, -26, 56 | 5 | 5.242 |
| Left Postcentral Gyrus | Postcentral_L | -48, -14, 58 | 5 | 5.323 |
| Left Paracentral Lobule | Paracentral_Lobule_L | -6, -30, 70 | 20 | 5.355 |
| Right Paracentral Lobule | Paracentral_Lobule_R | 10, -30, 68 | 16 | 5.611 |
| Right SMA | Supp_Motor_Area_R | 6, -20, 68 | 8 | 5.276 |
| Right Paracentral Lobule/Postcentral Gyrus | Paracentral_Lobule_R<br>Postcentral_R | 12, -34, 80 | 14 | 5.550 |
| Left Paracentral Lobule | Paracentral_Lobule_L<br>Precentral_L | -8, -20, 80 | 52 | 6.999 |
| ***Happy [Face_2 – Face_1] > Neutral [Face_2 – Face_1] (FDR-corrected cluster threshold, $p < 0.05$)*** |  |  |  |  |
| Right Limbic Lobe | Fusiform_R<br>ParaHippocampal_R<br>Lingual_R<br>Cerebellum_4_5_R<br>Hippocampus_R | 32, -36, -12 | 244 | 8.713 |
| Left Limbic Lobe | Fusiform_L<br>ParaHippocampal_L<br>Lingual_L<br>Cerebellum_4_5_L<br>Temporal_Inf_L | -30, -40, -10 | 197 | 7.027 |
| Left Retrosplenial Cortex | Calcarine_L<br>Cuneus_L<br>Precuneus_L<br>Lingual_L<br>Occipital_Sup_L<br>Vermis_4_5 | -18, -58, 14 | 150 | 6.375 |
| Right Retrosplenial Cortex | Precuneus_R<br>Calcarine_R<br>Lingual_R<br>Cuneus_R | 18, -54, 20 | 247 | 7.490 |
| Left Occipital-Parietal Cortex | Occipital_Mid_L<br>Angular_L<br>Parietal_Inf_L | -32, -84, 30 | 338 | 6.712 |
| Right Occipital-Temporal Cortex | Occipital_Mid_R<br>Angular_R<br>Occipital_Sup_R<br>Temporal_Mid_R | 36, -78, 38 | 273 | 6.092 |
| Right Precuneus | Precuneus_R<br>Parietal_Sup_R | 12, -70, 52 | 61 | 4.625 |
| Right Angular Gyrus | Angular_R | 42, -60, 50 | 9 | 4.350 |
| Right Superior Frontal Sulcus | Frontal_Sup_2_R<br>Frontal_Mid_2_R | 28, 12, 54 | 9 | 4.495 |
| Right Superior Frontal Sulcus | Frontal_Sup_2_R<br>Frontal_Mid_2_R | 32, 14, 62 | 19 | 4.424 |

**Figure 6.**
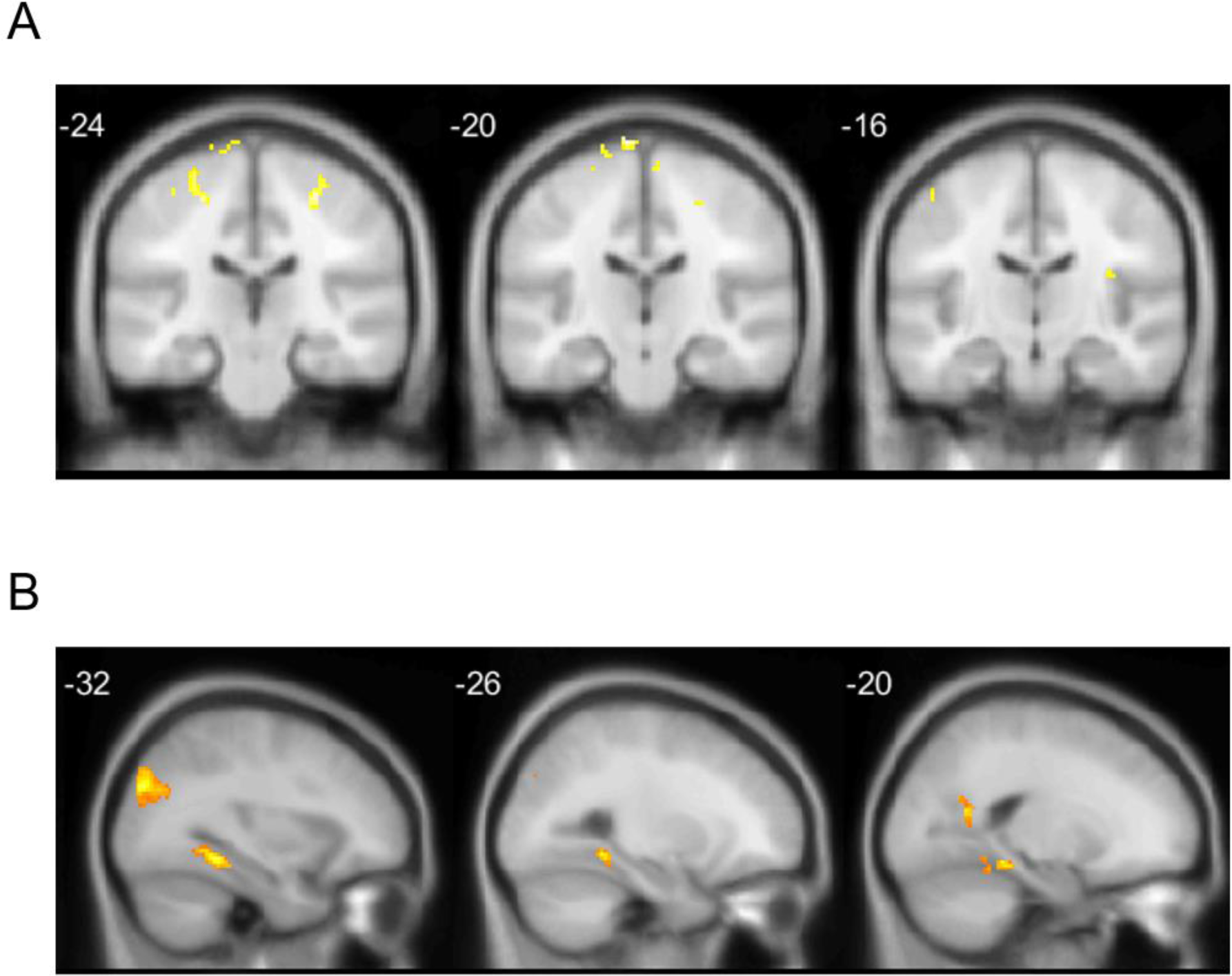
Interaction between neutral face processing and emotional conditions. (A) Brain regions showing the (Face_2 – Face_1) contrast in fearful condition minus the (Face_2 – Face_1) contrast in neutral condition. (B) Brain regions showing the (Face_2 – Face_1) contrast in happy condition minus the (Face_2 – Face_1) contrast in neutral condition (*p* < 0.05, FDR-corrected, cluster size > 5 voxels).

Similarly, analyzing the (Face_2 – Face_1) contrast in the happy condition against the neutral condition uncovered 10 clusters associated with happy scenes (Table 2 and Figure 6B). These regions, including the right hippocampus, bilateral PHC, bilateral AG, right MFG, right superior frontal gyrus (SFG), left IPL, right superior parietal gyrus (SPG), bilateral FG, bilateral retrosplenial cortex (RSC), left inferior temporal gyrus (ITG), left IPL, and right middle temporal gyrus (MTG), strongly indicate the presence of the Kuleshov effect in happy scenes. The activation patterns suggest heightened emotional memory engagement and increased attention to scene content in response to happiness, unveiling distinct brain regions contributing to the generation of the Kuleshov effect within happy scenes.

## 4. Discussion

While montage theory stands as a cornerstone in film studies, the early evidence from the original Kuleshov experiment remains ambiguous and lacks clarity (Pudovkin, 1970), leaving the existence of the Kuleshov effect an open question. Over the past three decades, despite replicated experiments suggesting the presence of the Kuleshov effect (Barratt et al., 2016; Calbi et al., 2017; Mullennix et al., 2019; Prince & Hensley, 1992), many of them relied on inauthentic film materials, such as static images. Consequently, their findings may not generalize to the existence of the Kuleshov effect in an authentic film scenario. This study aims to examine the Kuleshov effect using highly ecological films and explores whether film editing can influence viewers’ perception of emotion. Additionally, we delve into an exploration of the neural correlates underpinning the Kuleshov effect to further confirm its existence from a neurocinematic perspective (Hasson et al., 2008). These evidence from the current study can further support montage theory within realistic films.

In Experiment 1, we shot authentic films, including neutral faces and emotional scenes, and integrated them into a face-scene-face Kuleshov sequence (Figure 1). Before the Kuleshov experiment, we conducted a rating experiment to establish a baseline for neutral faces. During the Kuleshov experiment, participants were asked to rate the valence of the neutral face after watching the face-scene-face sequence, revealing that the emotional scene indeed affects emotion perception from a neutral face (Figure 3A). Specifically, fearful scenes prompt viewers to perceive negative emotion from a neutral face, while happy scenes prompt viewers to perceive positive emotion from a neutral face. Meanwhile, neutral scenes did not affect viewers’ perception of no expression from a neutral face. The ANOVA results showed a significant difference in valence between emotional and neutral conditions, suggesting that a viewer’s interpretation of an actor’s expression is significantly influenced by the emotional scene, leading to a context-dependent bias in emotional perception. Effectively controlling for the influence of prior knowledge about the Kuleshov effect ensures the reliability of our experimental results, as depicted in Figure S2. We conducted a similar Kuleshov experiment in fMRI in Experiment 2. In the behavioral results of Experiment 2, we found that the black screen inserted between emotional scenes and faces did not affect the observation of such a context-dependent bias in emotional perception (Figure 3C). Consistent with previous studies (Calbi et al., 2017; Mullennix et al., 2019), these behavioral results from Experiments 1 and 2 support the existence of the Kuleshov effect in an authentic film scenario. Notably, the Kuleshov sequence triggered higher arousal in the fearful condition compared with the neutral condition, consistent with prior research (Calbi et al., 2017). This observation indicates that the fearful condition not only initiates the context-dependent bias but also intensifies viewers’ emotional experiences. Additionally, we detected activation in multiple brain regions in the contrast between Face_2 and Face_1 in each condition (Figure 4) and in the contrast between Face_2 in the fearful or happy condition and the neutral condition (Figure 5). These are the neural correlates of the Kuleshov effect, and the activation of these regions supports the existence of its effect at the neural level.

### Enhancing ecological validity of the Kuleshov effect with authentic films

In contrast to previous behavioral studies, we adopted highly ecological films to ensure that participants perceived the scenes as integral components of a continuous film rather than isolated images or clips. This methodological choice, while pivotal, deviated from earlier investigations (Barratt et al., 2016; Calbi et al., 2017; Mullennix et al., 2019), which relied on facial expressions from the KDEF picture set and utilized zoom-in techniques to simulate dynamic faces from static images. However, this approach faced challenges in achieving realism, including subtle facial tremors induced by individuals remaining still, the natural occurrence of eye blinking during observations, and the static background (Ambadar et al., 2005; Ponech, 1997; Tan, 2018), making the handling of static images as real-life film clips difficult. Moreover, earlier investigations selected face and emotional scenes separately, rather than matching them as an integrated film. As the films closely mirror reality (Lu et al., 2016; Redcay & Moraczewski, 2020), adherence to continuity-editing rules aids viewers in perceiving the film as an emulation of the real world (Magliano & Zacks, 2011; T. J. Smith, 2012; T. J. Smith et al., 2012). Consequently, the utilization of isolated neutral faces and emotional scenes separately, without considering the coherence of the backgrounds, led to spatial discontinuities. This approach could potentially impact the examination of the existence of the Kuleshov effect.

To address these challenges, we captured neutral facial expressions against a blue background and subsequently replaced the background with images or videos, ensuring the seamless integration of neutral faces into the film’s reality (T. J. Smith et al., 2012). Furthermore, to mitigate potential influences from facial expressions, these faces were meticulously selected to maintain a neutral emotional state. In accordance with a previous study (Mullennix et al., 2019), we conducted a rating experiment, and the results for these neutral faces revealed a neutral valence (0.02 ± 0.13). Moreover, ensuring the consistency of neutral faces across emotional conditions is a prerequisite for observing Kuleshov effect bias. While some studies utilized neutral faces from the KDEF picture set, they did not assess the consistency of distinct groups (Barratt et al., 2016; Calbi et al., 2017). In the current study, the rating experiment we conducted indicated no significant differences among faces in various conditions (*F*_1,2_ = 1.23, *p* = 0.305).

### Confirming the existence of the Kuleshov effect through fMRI

Following exposure to the Kuleshov sequence, viewers attribute emotions to the neutral face (Figure 3), indicating a perceived new meaning for the second neutral face. To substantiate the existence of the Kuleshov effect, we explored the neural correlates associated with this new meaning. Contrasting activation in Face_2 and Face_1 revealed multiple activations (Figure 4, Table S1, S2, and S3), supporting the idea that the Kuleshov sequence indeed creates new meaning to the second neutral face, thus confirming the existence of the Kuleshov effect.

Specifically, our examination of the fearful condition unveiled activations in various brain regions, including the insula, cuneus, ACC, PCC, IPL, hippocampus, and OFC (Figure 4B). In contrast, both the happy condition (Figure 4C) and neutral condition (Figure 4D) displayed distinct activation patterns compared to the fearful condition, indicating the diversity in new meaning creation under different emotional contexts.

To comprehend why these regions exhibited activation upon exposure to neutral faces during the second viewing, we draw upon the notion that movie viewing encapsulates the intricacies of real-world interactions, enabling viewers to immerse themselves in the film (Redcay & Moraczewski, 2020). When scrutinizing the Kuleshov sequence, viewers engage in mental processes encompassing visual processing, theory of mind, and emotion generation. The activation of the cuneus in our findings may signifies visual processing of movie clips (Vanni et al., 2001). Simultaneously, the activation of the ACC, PCC, and IPL suggests that viewers employ theory of mind to interpret the film (Mahy et al., 2014; Mitchell & Phillips, 2015). Notably, the activation of the insula, a region associated with emotion processing, is noteworthy (Craig, 2011). The rationale behind emotion generation could be contextual processing (Zheng et al., 2022). Zheng et al. (Zheng et al., 2022) employing intracranial electroencephalogram recordings, investigated context-specific modulation among the amygdala, hippocampus, and OFC. They found that the OFC modulates the hippocampus and amygdala when perceiving emotions on a neutral face. The activation of the hippocampus and OFC in the current analysis aligns with Zheng’s study, supporting the idea that contextual processing may be the mechanism underlying the Kuleshov effect.

### Unveiling the neural correlates of the Kuleshov effect bias

In our exploration of the neural correlates of the Kuleshov effect bias using fMRI, we systematically examined Face_2 activations in fearful and happy conditions compared to neutral conditions (Figure 5). Notably, the contrast between Face_2 in fearful and neutral conditions revealed activations in key brain regions, including the precuneus, PHC, PCC, precentral gyrus, MTG, AG, and FG. These activations point to the involvement of these brain regions in contextual effects and emotional responses.

As highlighted by Mobbs et al. (Mobbs et al., 2006), contextual effects play a crucial role in shaping emotional attribution, aligning with previous studies on contextual information’s impact in various settings (Kumfor et al., 2018; Silveira et al., 2015). Kumfor’s exploration of emotional body language as contextual information revealed a correlation between abnormal contextual influence and reduced integrity in the right PHC and left precentral gyrus. Similarly, Silveira investigated contextual information using museum labels, demonstrating activations in the precuneus. The involvement of the PCC and AG, vital components of the default mode network (DMN), reinforces their role in contextual framing during the Kuleshov effect (Ranganath & Ritchey, 2012; V. Smith et al., 2021). The activations observed in the AG and FG during fearful conditions align with findings by Quinones Sanchez et al. (Quinones Sanchez et al., 2021), emphasizing heightened involvement in facial recognition and memory retrieval in fear-inducing circumstances. Additionally, the activation of the PCC, as noted in studies on emotional word processing (Maddock et al., 2003), further supports its role in evaluating emotional meanings.

Contrasting Face_2 between happy and neutral conditions revealed activations in the right cuneus, precuneus, and CAL. These findings support the existence of contextual effects in the happy condition (Silveira et al., 2015). The correlation between gray matter volume in the right precuneus and subjective happiness, as demonstrated by Sato (Sato et al., 2015), underscores the role of the precuneus in our activation results for perceiving happiness on neutral faces.

In our pursuit of understanding the Kuleshov effect bias, we conducted a comparative analysis by contrasting the (Face_2 – Face_1) activation in the fearful condition with a similar contrast in the neutral condition, revealing distinctive activation regions (Figure 6A). Notably, the insula emerged as a key region, supporting the concept that fearful conditions heighten the perception of negative emotion from a neutral face. Simultaneously, when examining the comparison between the happy and neutral contrasts (Figure 6B), discernible activation in the precuneus, hippocampus, and FG underscores their pivotal roles in contextual processing and its effects (Silveira et al., 2015; Zheng et al., 2022). This affirmation solidifies the understanding that the observed Kuleshov effect manifests as a contextual influence.

Collectively, our results significantly contribute to understanding the neural substrates of the Kuleshov effect, emphasizing the intricate interplay between contextual effects and emotional responses.

### Limitations and future directions

There are servals limitations in the current study. Firstly, it is crucial to acknowledge the potential impact of various factors on the observation of the Kuleshov effect, including viewer gender and film-watching experience (Ildirar & Ewing, 2018). While complete elimination of these influences remains challenging, we carefully balanced gender representation in our experiments and conducted a comprehensive questionnaire analysis to statistically assess participants’ film-watching frequency. The results, depicted in Figure S3, indicate that watching frequency did not significantly affect the observation of the Kuleshov effect. Secondly, the role of sound in creating an authentic film experience is essential, potentially contributing to the auditory Kuleshov effect (Baranowski & Hecht, 2017). Despite our intentional removal of all sound from the current film clips, aligning with a previous study (Calbi et al., 2017), the potential interaction between visual and auditory Kuleshov effects warrants exploration in future studies. Thirdly, while the amygdala’s involvement in fMRI studies of the Kuleshov effect has been documented (Mobbs et al., 2006), we did not detect its activation with FDR-correction in our study. This disparity may arise from paradigm differences, as Mobbs’ study employed a scene-face sequence with static images, while ours incorporated a face-scene-face sequence with authentic film clips. Considering the constraints of creating authentic films within budgetary limits, we captured a limited number of scenes and conducted a restricted number of trials in the current study. Future research should consider expanding the number of trials to further observe the activation patterns associated with the Kuleshov effect.

## 5. Conclusion

In this study, our objective was to investigate the existence of the Kuleshov effect using authentic films. We produced these film clips and conducted two experiments. The results from behavioral experiment indicate that viewers perceive emotions on neutral faces after watching the Kuleshov sequence, suggesting that film editing has the capacity to alter viewer perception by introducing new meanings. Through a comparative analysis of the valence of neutral faces across different emotional conditions, we identified a context-dependent bias in the Kuleshov effect. Specifically, viewers tend to perceive negative emotions on neutral faces after exposure to fearful film clips and positive emotions after watching happy film clips. Furthermore, our exploration extended to the neurobiological basis of the observed phenomena through fMRI experiments. The activation patterns in the insula, cuneus, precuneus, hippocampus, PCC, PHC, FG, and OFC suggest that the mechanism underlying the Kuleshov effect is contextual framing. These findings provide robust evidence supporting the existence of the Kuleshov effect at both behavioral and neural levels. Our study aligns with research utilizing the Kuleshov paradigm to investigate contextual framing (Calbi et al., 2017, 2019; Mobbs et al., 2006; Zheng et al., 2022), offering additional evidence that film editing can significantly influence viewer’s perception, evoking diverse emotional responses. Furthermore, our study contributes valuable insights to the understanding of film theory from a viewer’s perception viewpoint.

### Ethics statement

The studies involving human participants were reviewed and approved by Institutional Review Board of the State Key Laboratory of Cognitive Neuroscience and Learning at Beijing Normal University. The participants provided their written informed consent to participate in this study. Written informed consent was obtained from the individual(s) for the publication of any potentially identifiable images or data included in this article.

## Author contributions

**Zhengcao Cao**: Conceptualization, Methodology, Software, Formal analysis, Investigation, Visualization, Data Curation, Writing - original draft. **Yashu Wang**: Conceptualization, Methodology, Investigation, Writing - review & editing. **Liangyu Wu**: Resources. **Yapei Xie:** Methodology, Investigation, Software. **Zhichen Shi**: Investigation, Formal analysis. **Yiren Zhong:** Investigation. **Yiwen Wang**: Funding acquisition, Conceptualization, Supervision, Writing - review & editing.

## Funding

This study was financially sponsored through the Arts Project of 2019 National Social Science Fund of China (grant no.19BC041) and the Arts Project of 2023 National Social Science Fund of China (grant no.23ZD07). The authors thank them for financial support.

## Acknowledgement

We express our gratitude to Professor Yong He from IDG/McGovern Institute at BNU for the assistance in the data collection process. Special appreciation is extended to the film crew at Sichuan Film and Television University for their essential support in shooting films. The authors would also thank Ran Li, Xiang Xiao, Yanlin Zhu, Dayi Zhou and Xinya Huang for their support during the study.

## Conflict of Interest

The authors declare that the research was conducted in the absence of any commercial or financial relationships that could be construed as a potential conflict of interest.

## Notes

### Competing Interest Statement

The authors have declared no competing interest.

