## Supplementary Materials for "Revisiting the Kuleshov effect with authentic films: A behavioral and fMRI study"

#### Supplementary methods

1.1. Participants

Twelve healthy volunteers (6 females, age 26.75 ± 3.41 years) with normal or corrected-to-normal vision were recruited from the Beijing Normal University. Eligible participants indicated via a questionnaire that they did not have panic disorder. Participants majoring in film studies were also excluded.

1.2. Neutral faces rating procedure

In each trial, participants initiated the session by viewing a 2-second clip of a neutral face rendered in grayscale. Following the clip presentation, participants were tasked with evaluating the emotion of neutral faces with a scale. The first scale involved rating the valence of the neutral face on a scale from -4 to 4, where -4 denoted a negativity emotion, and 4 indicated a positivity emotion. The rating had a 3.5-second response time. Subsequently, a 2-second inter-stimuli interval (ISI) concluded the trial, and participants seamlessly progressed to the subsequent trial, completing a total of 30 trials. Stimuli delivery and response recording were controlled using PsychoPy 3.2 software (https://www.psychopy.org/), administered on a 14-inch laptop.

1.3. Emotional scenes rating procedure

In each trial, participants initiated the session by viewing a 4-second clip of an emotional scene rendered in grayscale. Following the clip presentation, participants were asked to judge the feeling of the emotional scene with a scale. The first scale involved rating the valence of the neutral face on a scale from -4 to 4, where -4 denoted a negativity feeling, and 4 indicated a positivity feeling. The rating had a 3.5-second response time. Subsequently, a 2-second ISI concluded the trial, and participants seamlessly progressed to the subsequent trial, completing a total of 30 trials. Stimuli delivery and response recording were controlled using PsychoPy 3.2 software, administered on a 14-inch laptop.

1.4. Data Analysis

For the neutral faces rating experiment, our hypothesis posits that all neutral faces exhibit similar valence, and the averaged valence of faces approaches zero. To test this, we calculated the average valence for faces under each type of emotional scene. Subsequently, a one-way analysis of variance (ANOVA) was conducted to confirm that there were no statistical differences in valence among the three emotional conditions.

In the emotional scenes rating experiment, our hypothesis posits that fearful scenes have negative valence, neutral scenes have neutral valence, and happy scenes have positive valence. To test this, we computed the average valence for all scenes under each type of emotional scene. Subsequently, a one-way analysis of variance (ANOVA) was conducted to confirm the statistical differences in valence for scenes among the three emotional conditions.

### Supplementary Results

**Figure S1. Examples of emotional scenes.**

**Figure S2. Behavioral results with subgroups of knowing Kuleshov effect in experiment 1.**

**Figure S3. Behavioral results with subgroups of watching frequency in experiment 1.**

**Table S1. fMRI Results: Face_2 minus Face_1 in fearful condition.**

**Table S2. fMRI Results: Face_2 minus Face_1 in happy condition.**

**Table S3. fMRI Results: Face_2 minus Face_1 in neutral condition.**

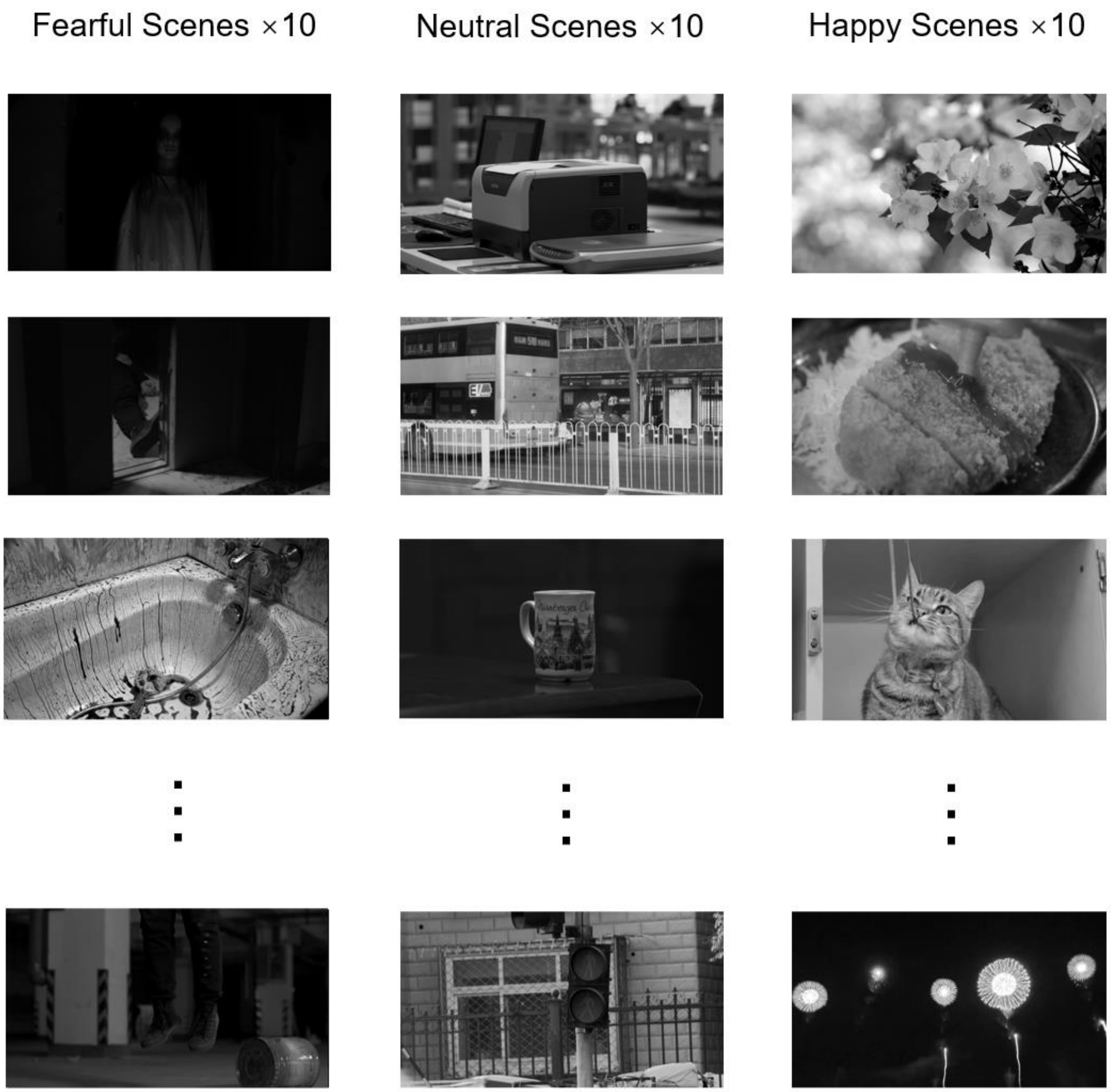


**Figure S1. Examples of emotional scenes.** These scenarios spanned various genres, including horror, documentary and comedy. Fearful scenes depicted chilling scenarios such as murder and ghosts. Neutral scenes featured commonplace settings like a bus stop, a printer, and a cup. Happy scenes showcased joyful images of flowers, food, and a cat.

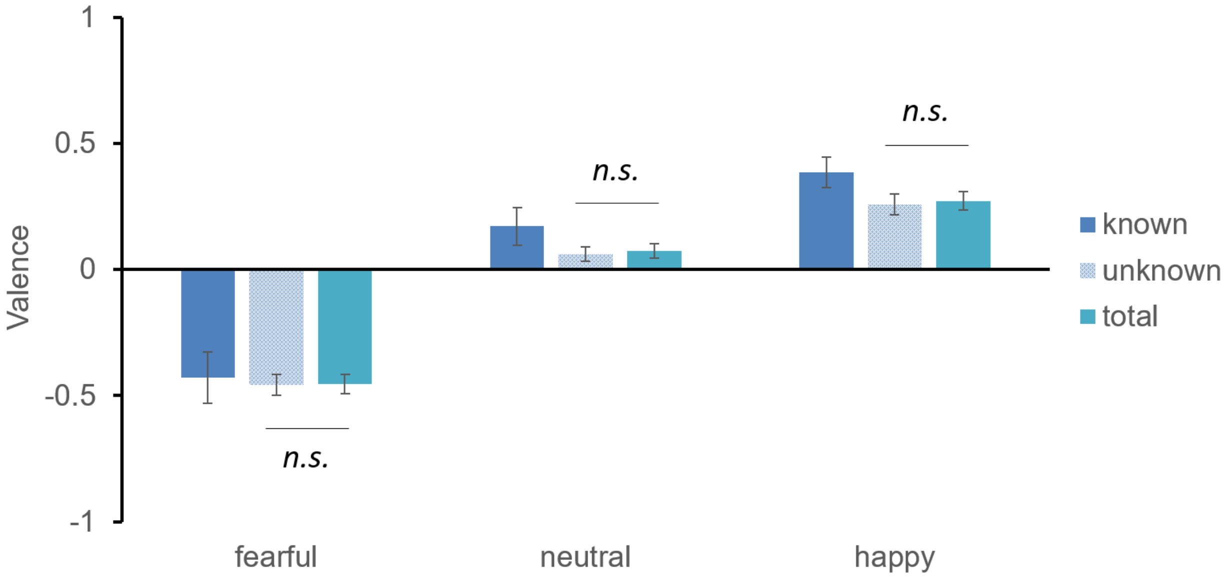


**Figure S2. Behavioral results with subgroups of knowing Kuleshov effect in experiment 1.** To identify and exclude participants with prior knowledge of the Kuleshov effect, we administered a knowledge test. Out of the total 59 participants, 7 were found to be familiar with the Kuleshov effect, while 52 were not. An ANOVA analysis conducted on valence ratings across these three groups revealed no significant difference between the group unaware of the Kuleshov effect and the total group. This suggests that the current validation of the Kuleshov effect is robust.

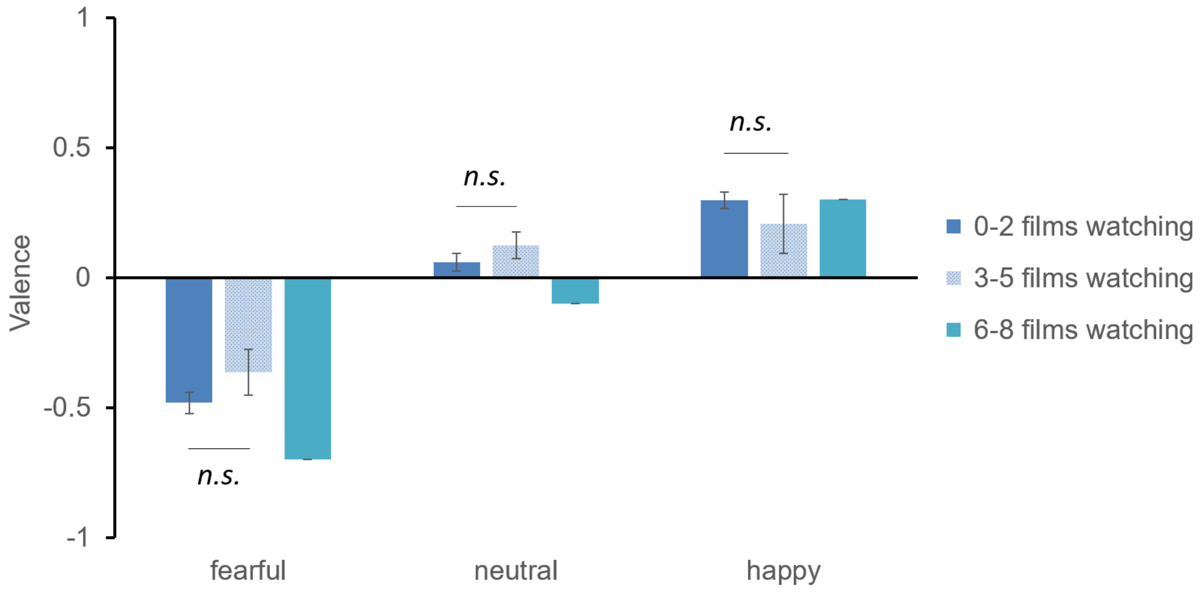


**Figure S3. Behavioral results with subgroups of watching frequency in experiment 1**. To examine whether film-watching frequency influences the observation of the Kuleshov effect, we conducted a subgroup comparison based on watching frequency. Out of the total 59 participants, 42 participants watched 0-2 films per week, 16 participants watched 3-5 films per week, and only 1 participant watched 6-8 films per week. Although there is a tendency for the middle-watching group (3-5 films per week) to have lower valence than the less-watching group (0-2 films per week), an ANOVA analysis conducted on valence ratings across these three groups revealed no significant difference between the less-watching group and the middle-watching group. This suggests that film-watching frequency does not affect the perception of the Kuleshov effect.

**Table S1. fMRI Results: Face_2 minus Face_1 in fearful condition.**

| Brain Region | AAL Atlas Labels | Peak Voxel Coordinate (MNI) | Cluster Size (KE) | T-score |
|---|---|---|---|---|
| ***Face_2 > Face_1*** *(FDR-corrected cluster threshold, p < 0.05)* | | | | |
| Cerebellum | Cerebellum_9_L<br>Cerebellum_9_R | 0, -46, -52 | 97 | 6.579 |
| Cerebellum | Cerebellum_8_R<br>Cerebellum_9_R<br>Cerebellum_7b_R | 26, -48, -52 | 396 | 6.737 |
| Cerebellum | Cerebellum_8_L<br>Cerebellum_9_L | -14, -62, -52 | 191 | 5.961 |
| Cerebellum | Cerebellum_6_L<br>Cerebellum_Crus2_L<br>Cerebellum_4_5_L<br>Vermis_4_5<br>Cerebellum_4_5_R<br>Cerebellum_6_R<br>Vermis_6<br>Cerebellum_Crus1_R<br>Cerebellum_Crus1_L<br>Cerebellum_Crus2_R<br>Cerebellum_7b_R<br>Cerebellum_7b_L<br>Vermis_3<br>Cerebellum_8_R<br>Vermis_7 | -44, -72, -42 | 2428 | 7.769 |
| Cerebellum | Cerebellum_Crus2_R<br>Cerebellum_Crus1_R | 36, -62, -42 | 12 | 3.204 |
| Cerebellum | Vermis_10 | 4, -46, -28 | 19 | 3.374 |
| Cerebellum | Cerebellum_6_R | 28, -40, -34 | 5 | 2.965 |
| Cerebellum | Cerebellum_8_R<br>Vermis_8<br>Cerebellum_Crus2_R | 6, -66, -32 | 7 | 3.174 |
| Cerebellum | Cerebellum_Crus1_L | -32, -66, -32 | 30 | 4.594 |

| | | | | |
|---|---|---|---|---|
| Right Inferior Temporal Gyrus | Temporal_Inf_R | 54, -16, -34 | 9 | 3.876 |
| Left Inferior Temporal Gyrus | Temporal_Inf_L | -52, -16, -34 | 5 | 3.599 |
| SMA/Augular Gyrus/Insula/STG/ACC /Hippocampus (bilaterally) | Postcentral_R<br>Postcentral_L<br>Precentral_R<br>Parietal_Inf_L<br>Parietal_Inf_R<br>Precentral_L<br>Angular_R<br>Frontal_Sup_2_R<br>Paracentral_Lobule_L<br>Supp_Motor_Area_R<br>Supp_Motor_Area_L<br>Parietal_Sup_L<br>Precuneus_R<br>Parietal_Sup_R<br>SupraMarginal_R<br>Rolandic_Oper_R<br>Rolandic_Oper_L<br>Cingulate_Mid_R<br>Paracentral_Lobule_R<br>Temporal_Inf_R<br>Angular_L<br>Temporal_Mid_R<br>Precuneus_L<br>Frontal_Sup_2_L<br>Frontal_Mid_2_R<br>Frontal_Inf_Oper_R<br>Insula_R<br>Temporal_Sup_L<br>Cingulate_Mid_L<br>SupraMarginal_L<br>Temporal_Sup_R | 18, -38, 18 | 20624 | 9.815 |

| | | | | |
|---|---|---|---|---|
| | Insula_L<br>Heschl_R<br>Heschl_L<br>Cuneus_R<br>Occipital_Mid_R<br>Caudate_L<br>Cingulate_Post_L<br>Thal_PuM_R<br>Frontal_Inf_Oper_L<br>Caudate_R<br>Occipital_Sup_R<br>Hippocampus_L<br>Occipital_Mid_L<br>Temporal_Pole_Sup_L<br>Cingulate_Post_R<br>Hippocampus_R<br>Temporal_Pole_Sup_R<br>Thal_PuM_L<br>Putamen_R<br>Thal_PuA_R<br>Occipital_Sup_L | | | |
| Cerebellum | Vermis_7<br>Cerebellum_Crus2_L | 0, -80, -26 | 20 | 3.785 |
| Left Temporal Lobe | Temporal_Mid_L<br>Temporal_Inf_L | -66, -26, -14 | 55 | 3.527 |
| Right Frontal Lobe | OFCant_R<br>Frontal_Mid_2_R<br>OFCmed_R | 28, 46, -14 | 111 | 5.299 |
| Right Temporal Lobe | Temporal_Inf_R | 60, -52, -16 | 7 | 3.581 |
| | OFCant_L<br>OFClat_L<br>Frontal_Mid_2_L | -26, 46, -14 | 26 | 4.119 |
| Right Temporal Lobe | Temporal_Mid_L | -54, -32, -8 | 12 | 3.323 |
| Left Frontal Lobe | Frontal_Sup_2_L | -26, 58, -10 | 7 | 3.201 |

| | | | | |
|---|---|---|---|---|
| Right Temporal Lobe | Temporal_Sup_R<br>Insula_R<br>Temporal_Pole_Sup_R | 48, -4, -4 | 50 | 4.475 |
| Right Frontal Lobe | Frontal_Mid_2_R<br>Frontal_Sup_2_R<br>Frontal_Inf_Tri_R | 42, 46, 22 | 738 | 6.363 |
| Left Temporal Lobe | Temporal_Sup_L | -40, -26, 2 | 13 | 3.846 |
| Right Frontal Lobe | Frontal_Sup_2_R | 26, 66, -2 | 19 | 3.755 |
| Right Temporal Lobe | Temporal_Sup_R | 60, -16, 6 | 8 | 3.734 |
| Right SFG | Frontal_Sup_2_R | 32, 64, 10 | 12 | 3.410 |
| Left STG | Temporal_Sup_L | -44, -34, 8 | 47 | 4.748 |
| Left IFG | Frontal_Inf_Tri_L<br>Frontal_Mid_2_L | -36, 42, 12 | 7 | 3.173 |
| Caudate | Caudate_R | 20, 24, 14 | 12 | 3.767 |
| Left Middle Frontal Sulcus | Frontal_Mid_2_L | -40, 48, 20 | 5 | 2.835 |
| Cuneus | Cuneus_L | 2, -88, 34 | 13 | 4.016 |
| Left Precentral Gyrus | Precentral_L | -60, 2, 26 | 25 | 3.572 |
| Right DMPFC | Frontal_Mid_2_R<br>Frontal_Sup_2_R | 26, 56, 32 | 45 | 3.652 |
| Right ACC | Cingulate_Mid_R<br>Frontal_Sup_Medial_L<br>ACC_sup_R | 2, 36, 32 | 32 | 3.509 |
| Left Middle Frontal Sulcus | Frontal_Mid_2_L | -38, 44, 30 | 6 | 2.856 |
| Precuneus | Cuneus_L<br>Precuneus_L | -8, -76, 32 | 19 | 3.492 |
| Precuneus | Precuneus_L | -10, -66, 40 | 33 | 4.189 |
| Left VMPFC | Frontal_Sup_Medial_L | 0, 28, 40 | 15 | 3.620 |
| Left MFG | Frontal_Mid_2_L | -32, 22, 50 | 5 | 2.976 |

**Table S2. fMRI Results: Face_2 minus Face_1 in happy condition.**

| Brain Region | AAL Atlas Labels | Peak Voxel Coordinate (MNI) | Cluster Size (KE) | T-score |
|---|---|---|---|---|
| ***Face_2 > Face_1*** *(FDR-corrected cluster threshold, $p < 0.05$)* | | | | |
| Cerebellum | Cerebellum_9_L | -10, -40, -56 | 19 | 5.364 |
| Cerebellum | Cerebellum_6_L<br>Cerebellum_4_5_L<br>Cerebellum_6_R<br>Cerebellum_Crus1_L<br>Cerebellum_Crus2_L<br>Vermis_4_5<br>Cerebellum_8_L<br>Cerebellum_4_5_R<br>Vermis_6<br>Cerebellum_8_R<br>Cerebellum_Crus1_R<br>Cerebellum_7b_L<br>Cerebellum_9_L<br>Cerebellum_Crus2_R<br>Cerebellum_7b_R<br>Vermis_7<br>Cerebellum_9_R<br>Lingual_L<br>Vermis_8<br>Cerebellum_3_R<br>Vermis_3<br>Lingual_R<br>Cerebellum_10_L | -26, -50, -26 | 3891 | 8.026 |
| Cerebellum | Cerebellum_8_R<br>Cerebellum_Crus2_R<br>Cerebellum_7b_R<br>Vermis_8<br>Cerebellum_9_R | 14, -62, -44 | 219 | 4.855 |
| Cerebellum | Vermis_8<br>Cerebellum_8_L | -2, -68, -34 | 52 | 5.447 |

| | | | | |
|---|---|---|---|---|
| | Vermis_7<br>Cerebellum_Crus2_L | | | |
| Right Temporal Lobe | Temporal_Inf_R<br>Temporal_Mid_R | 64, -24, -24 | 525 | 8.769 |
| Left Temporal Lobe | Temporal_Inf_L<br>Temporal_Mid_L | -56, -46, -14 | 57 | 4.084 |
| Cerebellum | Vermis_3<br>Cerebellum_3_L | 4, -40, -18 | 11 | 3.199 |
| SMA/Augular Gyrus/STG/ACC /Hippocampus (bilaterally) | Frontal_Mid_2_R<br>Frontal_Sup_2_R<br>Parietal_Inf_R<br>Parietal_Inf_L<br>Postcentral_R<br>Precuneus_R<br>Angular_R<br>SupraMarginal_R<br>Parietal_Sup_R<br>Supp_Motor_Area_R<br>Precentral_R<br>Cingulate_Mid_R<br>Parietal_Sup_L<br>Postcentral_L<br>ACC_sup_R<br>ACC_pre_R<br>Supp_Motor_Area_L<br>Paracentral_Lobule_L<br>ACC_sup_L<br>Precuneus_L<br>Cuneus_R<br>Frontal_Sup_Medial_L<br>Angular_L<br>Cingulate_Mid_L<br>ACC_pre_L<br>Frontal_Sup_Medial_R<br>Caudate_R | 18, -38, 16 | 14467 | 8.754 |

| | | | | |
|---|---|---|---|---|
| | Caudate_L<br>Occipital_Mid_R<br>Precentral_L<br>SupraMarginal_L<br>OFCant_R<br>Frontal_Sup_2_L<br>Occipital_Mid_L<br>Frontal_Med_Orb_R<br>Occipital_Sup_R<br>Paracentral_Lobule_R<br>Frontal_Inf_Tri_R<br>Frontal_Inf_Orb_2_R<br>OFCmed_R<br>Hippocampus_L<br>Calcarine_L<br>Thal_PuM_L<br>Thal_PuM_R<br>ACC_sub_R<br>Cuneus_L | | | |
| Left Frontal Lobe | Frontal_Sup_2_L<br>Frontal_Mid_2_L<br>OFCant_L | -28, 50, -10 | 116 | 4.676 |
| Right Insula | Insula_R<br>Frontal_Inf_Orb_2_R | 34, 20, -8 | 12 | 3.410 |
| Left Rolandic operculum/Insula/Heschl | Rolandic_Oper_L<br>Insula_L<br>SupraMarginal_L<br>Frontal_Inf_Oper_L<br>Temporal_Pole_Sup_L<br>Postcentral_L<br>Temporal_Sup_L<br>Precentral_L<br>Heschl_L | -44, 0, 12 | 503 | 5.912 |
| Right Rolandic operculum/Insula/Heschl | Rolandic_Oper_R<br>Insula_R | 50, -20, 20 | 274 | 5.619 |

| | | | | |
|---|---|---|---|---|
| | Heschl_R<br>SupraMarginal_R<br>Temporal_Sup_R<br>Postcentral_R | | | |
| Right Rolandic operculum/Insula/Heschl | Frontal_Inf_Oper_R<br>Rolandic_Oper_R<br>Insula_R<br>Precentral_R<br>Temporal_Sup_R<br>Temporal_Pole_Sup_R<br>Frontal_Inf_Tri_R<br>Putamen_R<br>Heschl_R | 40, 4, 2 | 840 | 6.420 |
| Precuneus | Precuneus_R<br>Calcarine_R | 32, -48, 0 | 89 | 7.648 |
| Left STG | Temporal_Sup_L | -42, -24, 0 | 6 | 3.217 |
| Putamen | Putamen_R | 30, -2, 2 | 7 | 3.651 |
| Left MFG/Insula | Frontal_Mid_2_L<br>Frontal_Inf_Tri_L<br>Insula_L | -20, 34, 10 | 85 | 4.571 |
| Left MFG | Frontal_Mid_2_L<br>Frontal_Inf_Tri_L | -40, 44, 16 | 53 | 3.434 |
| Calcarine | Calcarine_L | -16, -62, 12 | 5 | 2.969 |
| Putamen | Putamen_R | 28, -12, 10 | 7 | 3.517 |
| Insula | Insula_L | -28, 16, 12 | 8 | 3.156 |
| Left Thalamus | Thal_VL_L | -14, -6, 14 | 6 | 3.603 |
| Left STG | Temporal_Sup_L | -44, -40, 14 | 5 | 4.137 |
| Right Heschl | Heschl_R<br>Temporal_Sup_R | 34, -32, 14 | 17 | 3.820 |
| Right STG | Temporal_Sup_R<br>Rolandic_Oper_R | 46, -32, 18 | 5 | 3.234 |
| Right Thalamus | Thal_PuM_R | 6, -20, 16 | 7 | 3.232 |
| Precuneus/Cuneus | Precuneus_L<br>Cuneus_L<br>Calcarine_L | -12, -66, 28 | 332 | 4.594 |

| | | | | |
|---|---|---|---|---|
| | Occipital_Sup_L<br>Parietal_Sup_L | | | |
| Left Supramarginal Gyrus | SupraMarginal_L | -68, -26, 20 | 7 | 3.291 |
| ACC (bilaterally) | Cingulate_Mid_R<br>Cingulate_Mid_L<br>Precuneus_R<br>Cingulate_Post_R<br>Cingulate_Post_L<br>Paracentral_Lobule_R<br>Precuneus_L<br>Paracentral_Lobule_L | 8, -30, 42 | 661 | 6.6123 |
| Left Precentral Gyrus | Frontal_Inf_Oper_L<br>Precentral_L | -38, -2, 24 | 13 | 4.638 |
| Cuneus | Cuneus_L | 0, -88, 34 | 8 | 3.592 |
| Left ACC | Cingulate_Mid_L<br>ACC_sup_L | -8, 10, 34 | 40 | 3.612 |
| Right MFG | Frontal_Mid_2_R | 30, 26, 32 | 8 | 3.353 |
| Left MFG | Frontal_Mid_2_L | -36, 34, 42 | 5 | 3.066 |
| Right Precentral Gyrus | Precentral_R<br>Postcentral_R | 38, -20, 46 | 5 | 2.863 |
| Right SMA/ACC | Supp_Motor_Area_R<br>Cingulate_Mid_R | 8, -14, 52 | 9 | 4.154 |
| Left SMA | Supp_Motor_Area_L | -14, -2, 50 | 10 | 3.322 |
| Precuneus | Precuneus_L<br>Precuneus_R | -2, -72, 60 | 15 | 3.742 |
| Precuneus | Precuneus_L<br>Parietal_Sup_L | -12, -68, 64 | 5 | 2.837 |
| Precuneus | Precuneus_R<br>Paracentral_Lobule_R | 4, -46, 64 | 5 | 2.860 |
| Left Precentral Gyrus | Precentral_L<br>Postcentral_L | -22, -24, 76 | 17 | 3.377 |

**Table S3. fMRI Results: Face_2 minus Face_1 in neutral condition.**

| Brain Region | AAL Atlas Labels | Peak Voxel Coordinate (MNI) | Cluster Size (KE) | T-score |
|---|---|---|---|---|
| ***Face_2 > Face_1*** *(FDR-corrected cluster threshold, p < 0.05)* | | | | |
| Cerebellum | Cerebellum_8_R<br>Cerebellum_7b_R<br>Cerebellum_9_R<br>Vermis_8<br>Cerebellum_Crus2_R | 30, -50, -54 | 483 | 7.099 |
| Cerebellum | Cerebellum_6_L<br>Cerebellum_6_R<br>Cerebellum_Crus1_L<br>Cerebellum_4_5_L<br>Cerebellum_8_L<br>Vermis_4_5<br>Vermis_6<br>Cerebellum_Crus2_L<br>Cerebellum_4_5_R<br>Cerebellum_Crus1_R<br>Cerebellum_7b_L<br>Vermis_7<br>Cerebellum_9_L<br>Vermis_3<br>Vermis_8<br>Cerebellum_3_R | -4, -62, -18 | 3521 | 9.957 |
| Cerebellum | Cerebellum_9_L | -4, -42, -52 | 5 | 3.670 |
| Cerebellum | Cerebellum_Crus1_L<br>Cerebellum_Crus2_L | -46, -70, -36 | 9 | 4.263 |
| Cerebellum | Vermis_8<br>Cerebellum_8_R<br>Vermis_7 | 6, -66, -36 | 16 | 3.179 |
| Cerebellum | Cerebellum_8_L<br>Vermis_8 | -4, -66, -34 | 11 | 4.541 |
| Right Temporal Lobe | Temporal_Inf_R<br>Temporal_Mid_R | 62, -24, -22 | 329 | 7.421 |

| | | | | |
|---|---|---|---|---|
| Left Temporal Lobe | Temporal_Mid_L<br>Temporal_Inf_L | -62, -34, -16 | 14 | 3.514 |
| Left Temporal Lobe | Temporal_Inf_L | -52, -28, -18 | 6 | 2.901 |
| Right Insula/Putamen/Heschl /STG | Insula_R<br>Frontal_Inf_Oper_R<br>Rolandic_Oper_R<br>Putamen_R<br>Precentral_R<br>Temporal_Sup_R<br>Temporal_Pole_Sup_R<br>Heschl_R<br>Frontal_Inf_Tri_R<br>Pallidum_R | 36, 2, 8 | 1928 | 9.426 |
| Right OFC | Frontal_Mid_2_R<br>OFCant_R<br>Frontal_Sup_2_R<br>OFCmed_R | 34, 54, -12 | 151 | 5.005 |
| Left Insula/STG/Heschl /Hippocampus/Angular Gyrus/Thalamus | Postcentral_L<br>Parietal_Inf_L<br>SupraMarginal_L<br>Insula_L<br>Rolandic_Oper_L<br>Parietal_Sup_L<br>Temporal_Sup_L<br>Frontal_Inf_Oper_L<br>Precuneus_L<br>Precentral_L<br>Temporal_Pole_Sup_L<br>Caudate_L<br>Heschl_L<br>Putamen_L<br>Precuneus_R<br>Thal_VPL_L<br>Cingulate_Post_L<br>Hippocampus_L | -30, -56, 2 | 5972 | 7.862 |

| | | | | |
|---|---|---|---|---|
| | Thal_PuM_L<br>Angular_L<br>Frontal_Inf_Orb_2_L | | | |
| Right Insula | Insula_R<br>Frontal_Inf_Orb_2_R<br>OFCpost_R | 36, 20, -10 | 49 | 4.816 |
| Left SFG/OFC | Frontal_Sup_2_L<br>Frontal_Mid_2_L<br>OFCant_L | -22, 46, -2 | 93 | 4.418 |
| Left Insula/STG | Temporal_Sup_L<br>Insula_L | -38, -14, -6 | 16 | 3.428 |
| Right ACC/SFG/Hippocampus /Precuneus/STG /Precentral Gyrus | Postcentral_R<br>SupraMarginal_R<br>Parietal_Inf_R<br>Frontal_Mid_2_R<br>Frontal_Sup_2_R<br>Parietal_Sup_R<br>Cingulate_Mid_R<br>Supp_Motor_Area_R<br>ACC_sup_R<br>ACC_sup_L<br>Supp_Motor_Area_L<br>Cingulate_Mid_L<br>Angular_R<br>Precentral_R<br>Rolandic_Oper_R<br>ACC_pre_R<br>ACC_pre_L<br>Precuneus_R<br>Paracentral_Lobule_L<br>Temporal_Sup_R<br>Frontal_Sup_Medial_L<br>Frontal_Sup_2_L<br>Precentral_L<br>Frontal_Sup_Medial_R | 44, -36, 54 | 11662 | 8.370 |

| | | | | |
|---|---|---|---|---|
| | Caudate_R<br>Cuneus_R<br>Heschl_R<br>Cingulate_Post_L<br>Frontal_Med_Orb_R<br>Hippocampus_R<br>Cingulate_Post_R<br>ACC_sub_L<br>Occipital_Sup_R<br>Thal_PuM_R<br>Occipital_Mid_R | | | |
| Left Frontal Lobe | Frontal_Sup_2_L<br>Frontal_Mid_2_L | -32, 46, 0 | 6 | 3.408 |
| Left Heschl/Insual | Heschl_L<br>Insula_L<br>Temporal_Sup_L | -38, -24, 4 | 48 | 3.689 |
| Caudate | Caudate_R | 16, 24, 2 | 7 | 4.087 |
| Right SFG | Frontal_Sup_2_R | 24, 58, 4 | 25 | 3.479 |
| Putamen | Putamen_L | -26, -6, 2 | 9 | 3.511 |
| Right Heschl | Heschl_R<br>Temporal_Sup_R | 38, -26, 8 | 21 | 4.337 |
| Left Middle Frontal Sulcus | Frontal_Mid_2_L<br>Frontal_Inf_Tri_L | -36, 38, 22 | 230 | 5.499 |
| Right Heschl | Heschl_R | 34, -28, 10 | 7 | 3.049 |
| Caudate | Caudate_R | 24, 12, 16 | 41 | 5.263 |
| Caudate | Caudate_L | -18, -8, 18 | 5 | 2.957 |
| Left SFG | Frontal_Sup_2_L | -18, 48, 20 | 6 | 2.992 |
| Cuneus | Cuneus_L<br>Precuneus_L | -4, -74, 30 | 68 | 3.875 |
| Caudate | Caudate_R | 24, -4, 32 | 59 | 4.379 |
| Left Precentral Gyrus | Precentral_L<br>Frontal_Inf_Oper_L | -54, 4, 30 | 27 | 3.630 |
| Left SFG | Frontal_Sup_2_L | -26, 26, 30 | 7 | 3.579 |
| Left Angular Gyrus | Angular_L<br>Parietal_Inf_L | -36, -60, 40 | 73 | 3.508 |

| | | | | |
|---|---|---|---|---|
| | Parietal_Sup_L<br>Occipital_Mid_L | | | |
| Right MFG | Frontal_Mid_2_R | 38, 32, 38 | 101 | 4.150 |
| Right Posterior Cingulate Cortex/SMA | Cingulate_Mid_R<br>Paracentral_Lobule_R<br>Cingulate_Mid_L<br>Supp_Motor_Area_R<br>Precuneus_R | 10, -28, 42 | 366 | 5.813 |
| Right SFG | Frontal_Sup_2_R | 24, 34, 38 | 5 | 3.159 |
| Right SFG | Frontal_Sup_2_R | 20, 18, 38 | 7 | 4.193 |
| Left Precentral Gyrus | Precentral_L | -32, -10, 46 | 29 | 5.251 |
| Right SFG/SMA | Frontal_Sup_2_R<br>Supp_Motor_Area_R<br>Frontal_Sup_Medial_R | 16, 26, 56 | 251 | 5.264 |
| Right Precuneus | Parietal_Sup_R<br>Precuneus_R | 10, -68, 64 | 54 | 3.319 |
| Left Postcentral Gyrus | Postcentral_L | -22, -30, 56 | 5 | 3.231 |
| Right Paracentral Lobule | Paracentral_Lobule_R | 12, -30, 56 | 7 | 3.805 |
| Right Paracentral Lobule | Paracentral_Lobule_R<br>Precuneus_R | 4, -44, 64 | 6 | 3.182 |
| Left Paracentral Lobule | Paracentral_Lobule_L | 0, -22, 80 | 8 | 3.353 |
